# TMPRSS2 structure-phylogeny repositions Avoralstat for SARS-CoV-2 prophylaxis in mice

**DOI:** 10.1101/2021.01.04.425289

**Authors:** Young Joo Sun, Gabriel Velez, Dylan Parsons, Kun Li, Miguel Ortiz, Shaunik Sharma, Paul B. McCray, Alexander G. Bassuk, Vinit B. Mahajan

## Abstract

Drugs targeting host proteins can act prophylactically to reduce viral burden early in disease and limit morbidity, even with antivirals and vaccination. Transmembrane serine protease 2 (TMPRSS2) is a human protease required for SARS-CoV-2 viral entry and may represent such a target.^1–3^ We hypothesized drugs selected from proteins related by their tertiary structure, rather than their primary structure, were likely to interact with TMPRSS2. We created a structure-based phylogenetic computational tool 3DPhyloFold to systematically identify structurally similar serine proteases with known therapeutic inhibitors and demonstrated effective inhibition of SARS-CoV-2 infection *in vitro* and *in vivo*.^4,5^ Several candidate compounds, Avoralstat, PCI-27483, Antipain, and Soybean-Trypsin-Inhibitor, inhibited TMPRSS2 in biochemical and cell infection assays. Avoralstat, a clinically tested Kallikrein-related B1 inhibitor,^6^ inhibited SARS-CoV-2 entry and replication in human airway epithelial cells. In an *in vivo* proof of principle,^5^ Avoralstat significantly reduced lung tissue titers and mitigated weight-loss when administered prophylactically to SARS-CoV-2 susceptible mice indicating its potential to be repositioned for COVID-19 prophylaxis in humans.

## Main

The coronavirus disease 2019 (COVID-19), caused by SARS-coronavirus 2 (SARS-CoV-2), has spread globally causing over 1,740,000 deaths (WHO). Prophylactic and early-stage therapies are needed for high-risk populations. Even with vaccines, adjunctive therapies that mitigate viral entry or replication may attenuate disease severity and reduce viral spread by asymptomatic and early-stage patients. In response to the urgent need for therapeutics, there is investigation into repositioning existing drugs towards viral proteins (e.g., Remdesivir). An alternative strategy is to target human proteins utilized by viruses with small molecules. This approach can synergistically with vaccination and may be especially important for individuals where vaccination is contraindicated or deferred, to prevent viral transmission that may occur after vaccination, for front-line workers exposed to repeated high viral load, in countries where sophisticated vaccine delivery and storage is unavailable, and potentially for protection from viral mutation and other viruses using similar host mechanisms.

Transmembrane serine protease 2 (TMPRSS2) is a human serine protease that is a priming protease for the Spike glycoprotein found on the surface of all coronaviruses.^2,3^ TMPRSS2’s S1-peptidase domain is required for SARS-CoV-2 entry into host epithelial cells in the upper- and lower-respiratory tract,^1,7^ but it is not necessary for development or homeostasis in mice, making it an attractive drug target.^8^ It is yet to be determined whether TMPRSS2 inhibition mitigates SARS-CoV-2 infection *in vivo*. Camostat, a serine protease inhibitor originally developed for acute pancreatitis, inhibits TMPRSS2 *in vitro* and is in clinical trials (NCT04321096).^4,9^ However, Camostat’s plasma half-life is less than one minute, and its efficacy for COVID-19 is yet to be determined.^1,10,11^ Thus, identification of inhibitors targeting TMPRSS2 with improved pharmacokinetic properties remains important.

Conventional methods for identifying drug candidates typically employ high-throughput screening (HTS) or *in silico* screening using compound libraries previously tested in humans.^12^ *In silico* screening for TMPRSS2 has been challenging as there is no high-resolution TMPRSS2 molecular structure. Although HTS methods can rapidly screen thousands of compounds, there are certain limitations. HTS methods utilize only a few, generalized experimental parameters with technical limitations, such as narrow dose range and experimental conditions, which may not account for the unique features of each compound. This can lead to false positives and negatives. While false positives are filtered out in subsequent experiments, false negatives may overlook valuable compounds. Because HTS uses shotgun rather than hypothesis-driven approaches, it may be difficult to ascertain the mechanism-of-action, and this may slow the downstream development of candidate drugs into human therapies. Hypothesis-driven screening methods, utilizing protein structures and a limited number of compounds, remains a valuable and complementary strategy for drug-repositioning. One approach to rational-based drug repositioning is to identify proteins with preexisting drugs that are similar to the target protein.

### In silico drug repositioning by 3DPhyloFold

To identify drug repositioning candidates, we created a computational/hypothesis-driven drug repurposing method called 3DPhyloFold that identifies structurally similar proteins to rationally select candidate inhibitors.^10^ A comprehensive phylogenetic analysis of 600 S1-peptidases sequentially related to the TMPRSS2 S1-peptidase domain (TMPRSS2-S1P) showed TMPRSS2-S1P clustered closely to canonical TMPRSS family members, like Hepsin, as well as proteases outside of the TMPRSS subfamily: Coagulation Factor XI and Kallikrein-related B1 (KLKB1; **Extended Data Fig. 1a, b**). TMPRSS2-S1P was closest to Hepsin, and a homology-based model was generated (**Extended Data Fig. 1c**). Next, 3DPhyloFold determined the 3D relationship of TMPRSS2-S1P to other S1-peptidase structures (**Fig. 1a**; **Supplementary Table 2**). Using structural quality metrics (see **Methods**), 74 S1-peptidases and TMPRSS2-S1P were aligned by conventional sequence phylogenetic analysis. TMPRSS2-S1P clustered closely with KLKB1, Factor XI, and Complement Factor I (CFAI) (**Fig. 1b**). The Kallikrein- and Trypsin-like clades clustered further away, indicating TMPRSS2-S1P was sequentially divergent (**Fig. 1b**). In 3DPhyloFold, pairwise structural comparisons of the representative tertiary structures were used to calculate a structural dissimilarity matrix (SDM) based on the root mean square deviation between protein alpha-carbons (C;α RMSD; **Fig. 1a**). A structure-based phylogenetic tree was then generated from the SDM (**Fig. 1c**). Clustering of the structure-based tree was distinct to that of the sequence-based tree. Proteins close in the primary sequence analysis (e.g., CFAI and CTRB1) were much farther away in the 3DPhyloFold structure-tree (**Extended Data Fig. 2g**). Although distant in the sequence-based tree, the Trypsin-like clade and Factor VII moved much closer to TMPRSS2-S1P in the structure-based tree (**Fig. 1c**). This suggested that, while divergent in sequence, TMPRSS2-S1P adopts a three-dimensional fold closer to Trypsin and Factor VII. We prioritized the six S1-peptidases with the highest structural similarity to TMPRSS2-S1P: Hepsin, Acrosin, Trypsin, Factor VII, Factor XI, and KLKB1.

**Fig. 1.**
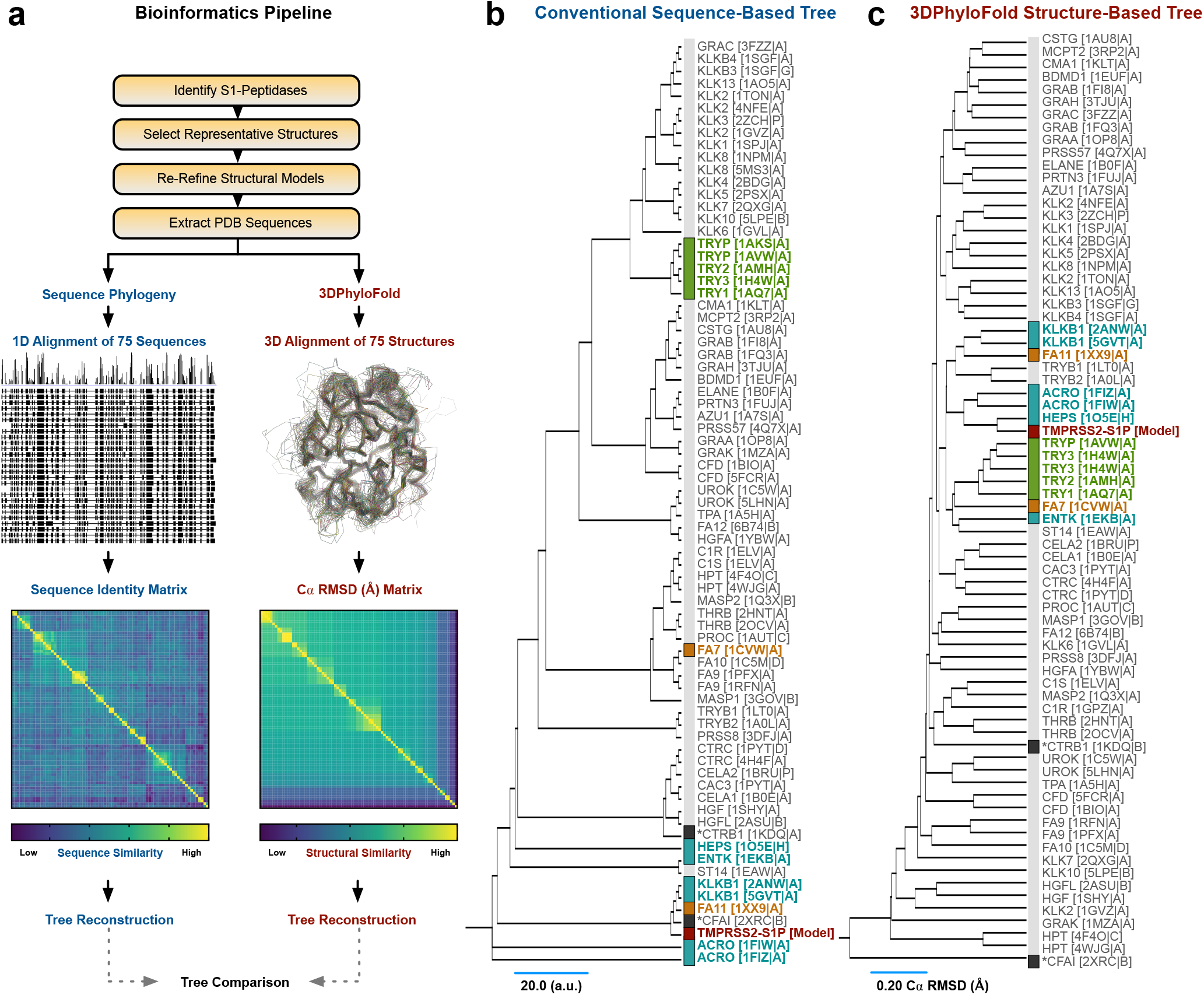
Structure-based phylogenetic analysis identifies sequentially divergent serine proteases with similar folds to TMPRSS2. **(a)**Bioinformatics workflow. **(b)**Sequence-based phylogenetic tree of 75 representative peptidase domains with the main clusters highlighted in different colors. Branches are labeled according to the corresponding structures and PDBIDs. **(c)**Structure-based phylogenetic tree of mammalian S1-peptidase domain structures. Evolutionary distance was inferred using the UPGMA method. Branch lengths correspond to the Cα RMSD (Å) of the pairwise-structural-alignments calculated in 3DPhyloFold. The proteases with the highest structural similarity to TMPRSS2-S1P were Hepsin, Acrosin, Trypsin, Factor VII, Factor XI, and KLKB1.

Using these six proteases, we sought known small-molecules and peptidomimetic inhibitors containing a guanidine, or structurally related groups (see **Methods**), since S1-peptidases are inhibited by compounds containing a 4-amindinobenzylamide moiety, a key specificity feature of their substrates where the first N-terminal residue at the cleavage site (P1) forms a strong interaction with an aspartate in the corresponding S1 subpocket (**Fig. 2a**).^13^ This search curated ninety experimental compounds and four small molecules previously tested in human clinical trials, which docked well to TMPRSS2-S1P (**Fig. 2b, c; Supplementary Table 3-4; Extended Data Fig. 3**). In addition, 3DPhyloFold analysis revealed a natural Trypsin-inhibiting protein based on the structure of porcine-Trypsin with Soybean-Trypsin-Inhibitor (SBTI; PDBID 1AVW; **Fig. 2c**). Since the porcine Trypsin binding pocket was similar to that of TMPRSS2-S1P (~68% sequence identity), we modeled the TMPRSS2-S1P/SBTI complex and identified a conserved inhibitory motif (PYRIRF), with favorable docking suggesting SBTI might bind and inhibit TMPRSS2-S1P (**Extended Data Fig. 4; Supplementary Table 4-5**).

**Fig. 2.**
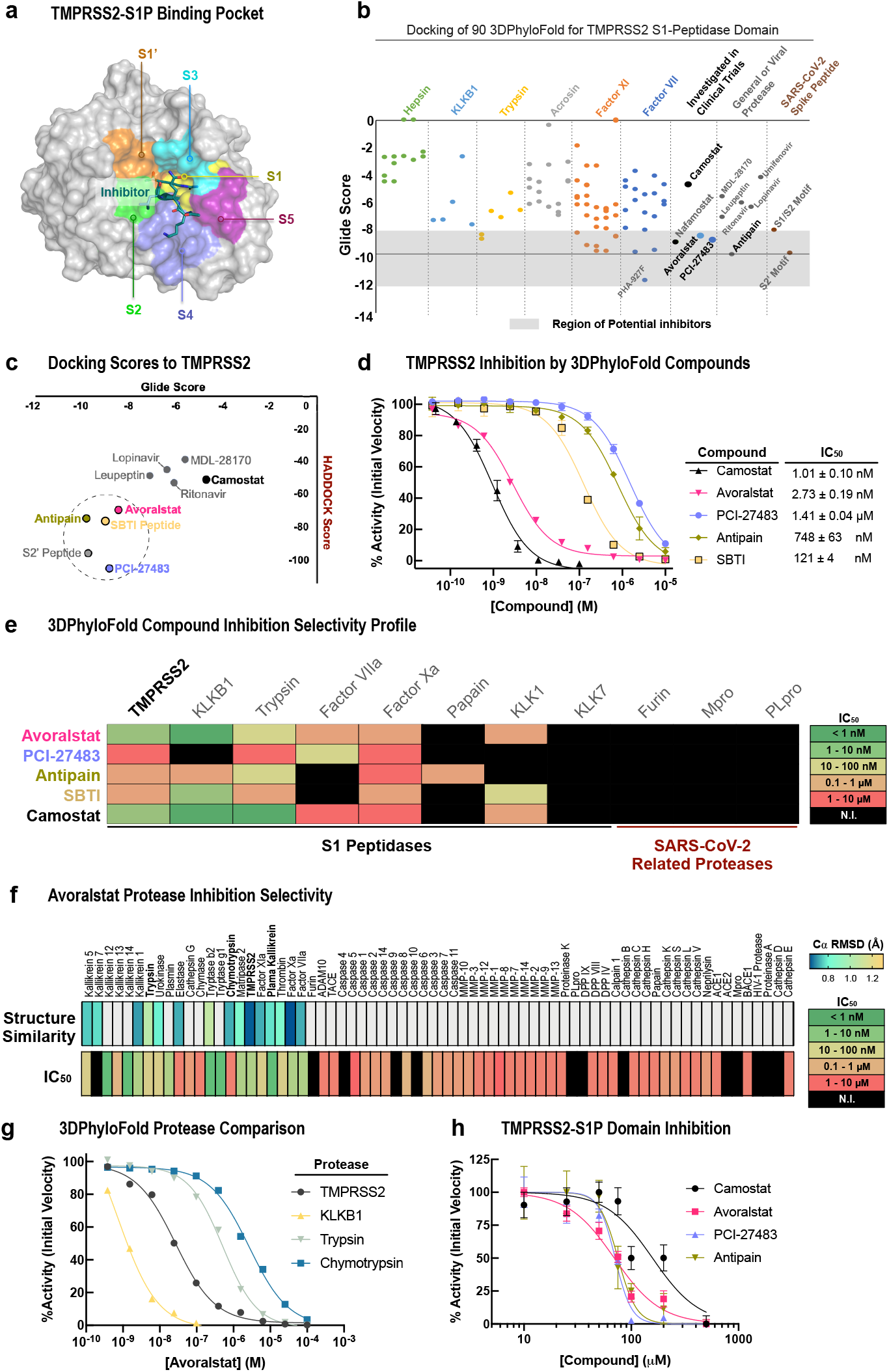
Compounds derived from 3DPhyloFold *in silico* analysis inhibit TMPRSS2 activity *in vitro*. **(a)** TMPRSS2-S1P structural model ligand-binding sub-pockets (S1’, S1, S2, S3, S4, S5). **(b)** Docking scores of compounds curated from 3DPhyloFold and **(c)** correlation between algorithms. Potential inhibitors clustered around the natural S2’-peptide motif. (**d)** TMPRSS2 inhibition by 3DPhyloFold compounds. **(e)** IC_50_ value selectivity profile against S1- and SARS-CoV-2-related proteases. **(f)** Avoralstat selectivity against 70 proteases. Top row - Cα-RMSD of pairwise-structural-alignments to TMPRSS2. **(g)** Protease inhibition by Avoralstat correlates with 3DPhyloFold prediction. **(h)** TMPRSS2-S1P domain inhibition. IC_50_ data represent mean ± SEM; n=3.

### Biochemical evaluation of 3DPhyloFold inhibitors

We focused on the inhibitory potential of human drugs available for repositioning, including Avoralstat, PCI-27483, and Antipain, along with SBTI. A biochemical inhibition assay using the extracellular compartment of purified recombinant TMPRSS2 (residues 106 – 492) was utilized to test compounds inhibition.^4^ The rank order potency against TMPRSS2 was Avoralstat (IC_50_ = 2.73 ± 0.19 nM), SBTI (IC_50_ = 121 ± 4 nM), Antipain (IC_50_ = 748 ± 63 nM), and PCI-27483 (IC_50_ = 1.41 ± 0.04 μM; **Fig. 2d**). Inhibition by Avoralstat was as potent as Camostat (IC_50_ = 1.01 ± 0.10 nM) which targets TMRPSS2 and is currently under clinical investigation for SARS-CoV-2 treatment.^4^ We further explored the selectivity profile of the four 3DPhyloFold inhibitors and Camostat as a positive control by testing them against six S1-peptidases identified in 3DPhyloFold (i.e., KLKB1, Trypsin, Factor VIIa, Factor Xa, KLK1, and KLK7), three other proteases involved in SARS-CoV-2 infection (i.e., Furin, Mpro, and PLpro), and a negative control Papain. As expected, each 3DPhyloFold compound displayed potent inhibition of its original target-proteases, and there was no inhibition of non-S1-proteases. Strikingly, Avoralstat was more than 18-fold selective towards TMPRSS2 than other S1-protease (**Fig. 2e; Supplementary Table 6**). Camostat was not as selective as Avoralstat. We further characterized Avoralstat specificity by expanding the protease screen to include additional 60 structurally distant proteases, including MMPs, Caspases, Cathepsins, and cysteine-or aspartyl-proteases.^1,14–16^ Avoralstat displayed potent inhibition of other S1-proteases consistent with their proximity to TMPRSS2 in the 3DPhyloFold tree, while displaying no inhibition of non-S1-proteases (**Fig 2f; Supplementary Table 7**), suggesting inhibition was specific and not due to protein aggregation effects. Notably, Avoralstat inhibited several proteins that were structurally similar to TMPRSS2, including Factor VIIa and Tryptase b2, despite being distant in primary sequence (**Fig. 2f**). Conversely, Avoralstat was less effective at inhibiting proteases that clustered further from TMPRSS2 on the 3DPhyloFold tree: Chymotrypsin (IC_50_ = >1 μM) and Elastase, (IC_50_ = >1 μM; **Fig. 2g**), despite their proximity in the sequence phylogenetic tree. To further confirm that the compounds target the protease domain of TMPRSS2, we tested inhibition using recombinant S1P domain (residues 252 – 489) and found similar inhibition trends (**Fig. 2h**). Taken together, these results suggested Avoralstat was highly selective for TMPRSS2, consistent with the predictions by structural phylogenetic analysis.

### Cellular evaluation of 3DPhyloFold inhibitors

Inhibition of full-length TMPRSS2 (TMPRSS2-FL) proteolytic activity was then tested in cells. TMPRSS2-FL contains an autoproteolysis motif (residues 252-257), which is subject to cleavage and can be used to probe the activity of TMPRSS2 in cells.^17^ Cells were transfected with either wild-type (WT) or loss of function TMPRSS2-S441A mutant (**Fig. 3a; Extended Data Fig. 6**). Compared to the inactive S441A mutant, TMPRSS2-WT showed reduced signal by immunoblot as previously reported (**Fig. 3a**).^17^ Inhibitor treatment prevented TMPRSS2-FL autoproteolysis and significantly increased the TMPRSS2-FL band intensity **(Fig. 3a**).

**Fig. 3.**
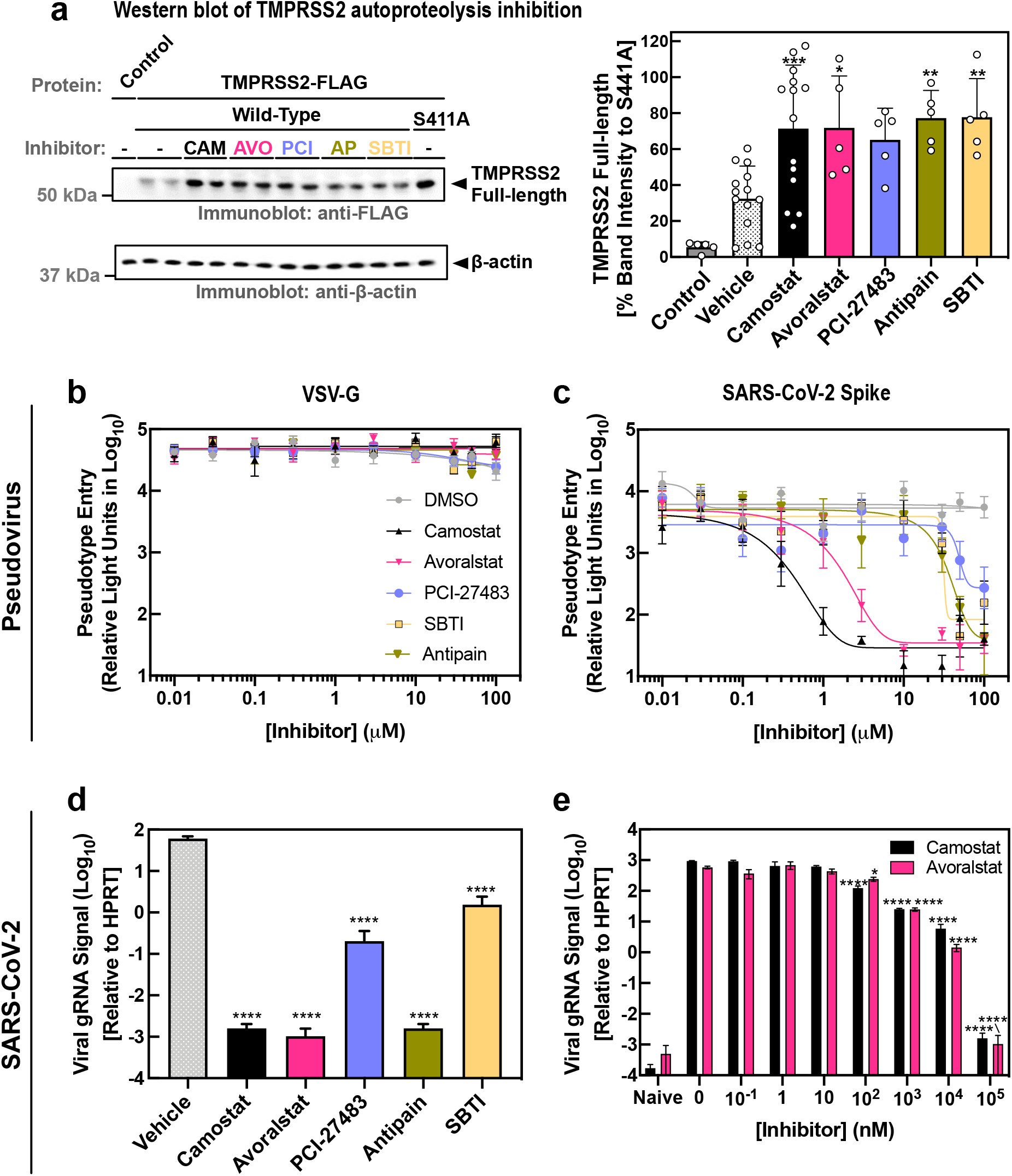
Avoralstat inhibits viral entry directed by SARS-CoV-2 spike proteins. **(a)** HEK-cells treated with Camostat, Avoralstat, PCI-27483, Antipain, or SBTI 2-hr before TMPRSS2 transfection. TMPRSS2 reduced autoproteolysis (increased TMPRSS2-FL signal; *p<0.0332, **p<0.0021, ***p<0.0002). Calu-3-cells were treated with compounds and inoculated with pseudovirions harboring **(b)** VSV-G or **(c)** SARS-CoV-2-spike-protein. Calu-3 cells were treated with **(d)** 100 μM or **(e)** indicated concentrations of each compound, then incubated with SARS-CoV-2. Viral gRNA was measured after 24-hrs. Data represent mean ± SEM; n=3. **(d)** Compounds reduced viral signal at 100 μM (****p<0.0001). **(e)** Viral signal was reduced beginning from 100 nM (*p<0.0332, ****p<0.0001).

To test whether the compounds specifically inhibit the molecular entry-pathway, transduction and infection assays were performed using vesicular stomatitis virus (VSV)-based pseudovirions bearing the SARS-CoV-2 Spike glycoprotein and a firefly luciferase reporter. Human Calu-3 2B4 airway cells were incubated with Camostat, Avoralstat, PCI-27483, Antipain, and SBTI. Pseudovirions harboring the pantropic VSV glycoprotein (VSV-G) served as controls since they transduce cells independent of TMPRSS2.^18^ Indeed, no compounds were toxic to cells and none prevented VSV-G pseudovirus entry, since the luciferase signal remained constant (**Fig. 3b**). Camostat inhibited SARS-CoV-2 pseudovirus entry (EC_50_ = 0.7 ± 0.2 μM), and Avoralstat displayed similar inhibition (EC_50_ = 2.8 ± 0.7 μM). PCI-27483, Antipain, and SBTI displayed modest inhibition but were too weak to determine reliable EC_50_ values (**Fig. 3c**).

Next, inhibition of authentic SARS-CoV-2 was tested in Calu-3 2B4 cells by measuring viral genomes. Camostat, Avoralstat, and Antipain significantly reduced SARS-CoV-2 replication (the amount of nucleocapsid gene [viral RNA] compared to vehicle, respectively; p<0.0001). PCI-27483 and SBTI showed less inhibition (**Fig. 3d**). A dose-response of Camostat and Avoralstat displayed significant reduction in SARS-CoV-2 infection beginning at 100 nM. Camostat and Avoralstat showed more than a ten-fold decrease in viral RNA signal with a 1 μM dose (**Fig. 3e**). SARS-CoV-2 showed more sensitivity to Avoralstat and Camostat than MERS-CoV, another coronavirus that also uses TMPRSS2 to facilitate entry (**Extended Data Fig. 7**).^9^

### Avoralstat inhibits SARS-CoV-2 entry in vivo

No therapy targeting host sensitizing proteases has been validated in an *in vivo* model of COVID-19. There is no known viral infection dose or animal model that fully recapitulates human disease, so the critical *in vivo* measure for testing prophylactic efficacy is the reduction of viral load. Using a mouse model of SARS-CoV-2 lung infection (Ad5-hACE2 transduced wild-type BALB/c mice^5^), we compared the efficacy of Avoralstat and Camostat in modifying SARS-CoV-2 infection. Cohorts of mice were infected intranasally with either 3 × 10^3^ or 1 × 10^5^ PFU of SARS-CoV-2, respectively. Mice were treated with Avoralstat, Camostat (30 mg/kg intraperitoneal injection), or vehicle (DMSO). Lungs were harvested 1 day after infection and viral titers measured by plaque assay. Both Avoralstat and Camostat significantly reduced the lung tissue titers in both cohorts (**Fig. 4a-b**). In a third cohort of mice, Avoralstat or Camostat was administered 4 hours prior and 4 hours after a 1 x 10^5^ PFU of SARS-CoV-2 intranasal challenge. Mice were given twice daily drug doses for three days post infection (dpi). Lungs harvested at 5-dpi showed both drugs significantly reduced the viral titers. Strikingly, the lung tissue virus titers were below the limit of detection in 3 of 4 Avoralstat-treated mice (**Fig. 4c**). Changes in weight, indicating the severity of illness, was monitored. Beginning at 4-dpi, there was significant weight loss in the vehicle- and Camostat-treated mice, while the weight of the Avoralstat-treated mice remained relatively constant suggesting a significant protective effect (**Fig. 4d**). Even though there was significant weight-loss in Camostat group, an Avoralstat therapeutic effect was observed later at 7-dpi compared to the vehicle-treated groups (**Fig. 4d**). In a fourth cohort of mice, a biological dose-response was strongly supported after we further increased the SARS-CoV-2 challenge dose to 1 × 10^6^ PFU. Avoralstat or Camostat were administered 4 hours prior and 4 hours after a SARS-CoV-2 intranasal challenge. Mice were then given two drug doses daily for 3-dpi. At the higher challenge dose, an early viral titer reduction was not observed as seen in lower titers (i.e., 3 × 10^3^ or 1 × 10^5^ PFU). Yet a significant decrease of viral titer was observed at 4-dpi for both Avoralstat- and Camostat-treated groups (**Fig. 4e**). Moreover, Avoralstat still showed a significant weight rescue effect beginning from 7-dpi while Camostat did not show any rescue effect compared to the vehicle-treated group (**Fig. 4f**). Thus, the inhibitory effect of Avoralstat observed in biochemical and cell assays, extended to prophylactic treatment of mice infected with escalating doses of SARS-CoV-2.

**Fig. 4.**
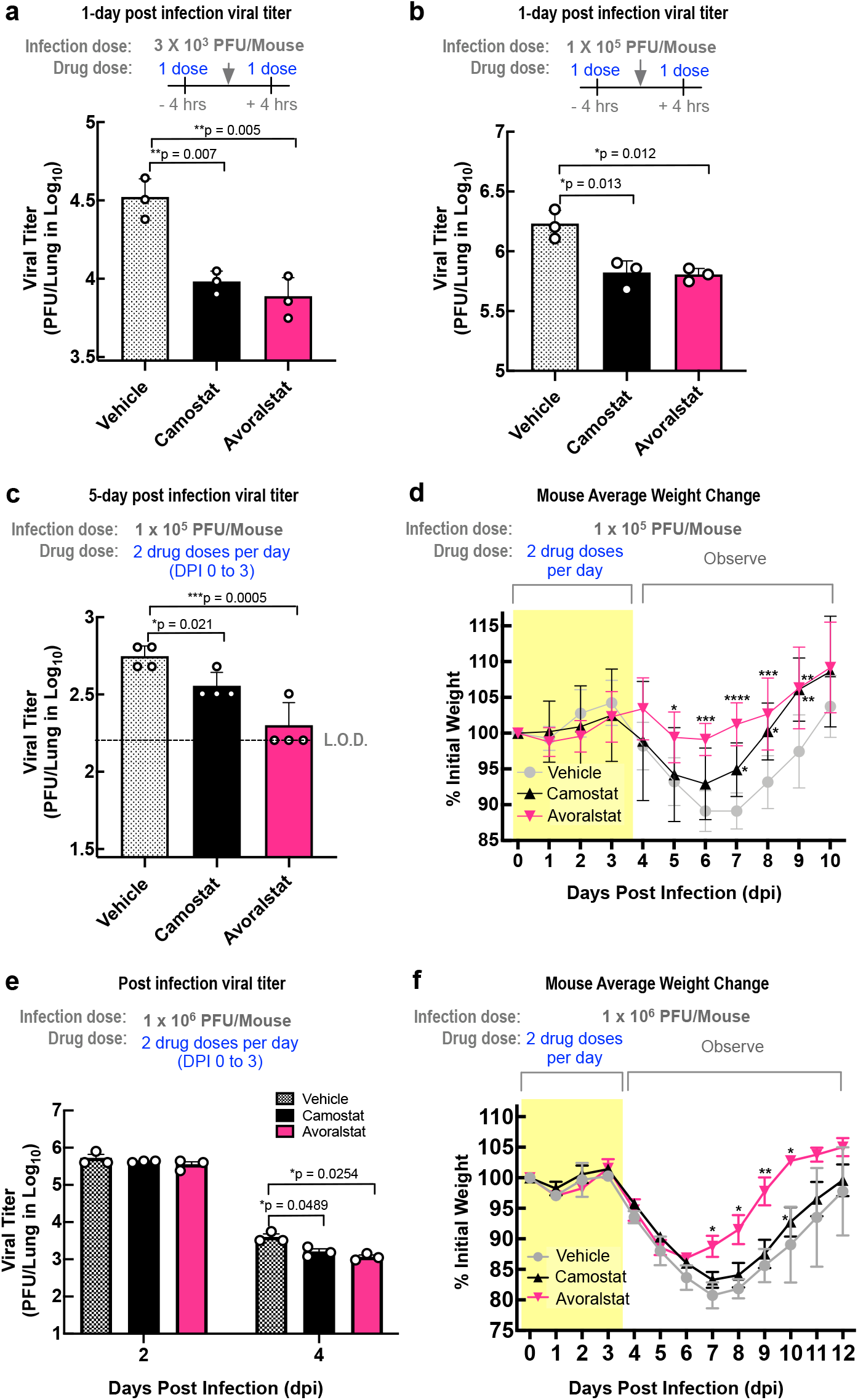
Avoralstat reduces SARS-CoV-2 infection in mice. **(a-c)** Ad5-hACE2-transduced BALB/c mice treated with Avoralstat, Camostat, or DMSO 4-hrs before and after SARS-CoV-2 viral challenge showed significantly reduced lung viral titer. **(d)** Avoralstat protected mice from weight-loss better than Camostat. **(e)** Avoralstat significantly reduced lung viral titer. **(f)** At the highest dose of SARS-CoV-2, Avoralstat protected mice from weight-loss. **(f)** Avoralstat and Camostat significantly lowered viral titer, n=6. (*p<0.0332, **p<0.0021, ***p<0.0002; mean ± SEM; L.O.D., limit of detection).

Drug repositioning is an important strategy to address human disease at a faster pace than conventional drug development, especially in the setting of a global viral pandemic. Avoralstat, a clinically tested oral KLKB1 inhibitor evaluated for the treatment of hereditary angioedema, successfully inhibited SARS-CoV-2 infection and illness in mice. Avoralstat is orally bioavailable, which could facilitate prophylactic administration to people at high risk for COVID-19, particularly where specialized transport, cold storage, and medically skilled delivery staff are not available. Avoralstat possess a favorable plasma half-life of 12-31 hours, compared to the short half-life of Camostat due to an easily cleavable ester bond, giving it a terminal half-life of roughly 1 hour. ^19,20,6,21^ It is possible that the observed efficacy of Avoralstat compared to Camostat in our *in vivo* study is due to its longer plasma half-life. Avoralstat has relatively minor and manageable side effects. No grade 3 adverse events were observed in phase 1 through phase 3 clinical trials for Avoralstat and no serious adverse events were more prevalent than those treated compared to placebo groups ^6,21^.The doses we tested in mice were a significantly lower dose than previously administered to humans in clinical trials, ^22^ suggesting an appropriate dose of Avoralstat for treating COVID-19 may be readily achievable with reasonable safety.

The application of a targeted structure-based phylogeny approach allowed us to identify and rationally prioritize several candidate TMPRSS2 inhibitors not considered by other drug repositioning strategies: 3DPhyloFold pointed to closely related proteins missed by primary-sequence comparisons, supporting a mechanism-based/hypothesis-driven selection of curated inhibitor candidates. Many of the small molecules tested in this study could be further developed for alternative routes of administration and for more potency and selectivity against TMPRSS2. The SBTI protein might serve as a cheap and natural source inhibitor for TMPRSS2, since it has been widely used in biomedical research.^23^ Interestingly, Avoralstat and PCI-27483 were both represented in high throughput screens but may have been missed due to the lack of sufficient testing conditions (e.g. dosage or cell line).^24^

Our *in vivo* studies underscore that targeting TMPRSS2 is a tenable strategy for COVID-19 treatment. A reduction of viral load achieved by an alternative mechanism to that of vaccination could act synergistically to reduce illness and transmission. In addition, TMPRSS2 is implicated in the cleavage of the envelope-glycoproteins of many other viruses, including SARS-CoV, MERS-CoV, HCoV-229E, HCoV-OC43, HCoV-HKU1, and HCoV-NL63; Influenza viruses; Parainfluenza viruses; and human Metapneumovirus.^10^ Thus, targeting this host machinery could be applied as a long-term strategy for future zoonotic coronaviruses and other respiratory viruses. This may be especially important if targeting viral proteins are only partially effective, natural infection does not confer long-lasting immunity, and combination therapies are needed to reduce the likelihood of resistance.^25^

## End Notes Acknowledgments

The following reagent was deposited by the Centers for Disease Control and Prevention and obtained through BEI Resources, NIAID, NIH: SARS-Related Coronavirus 2, Isolate USA-WA1/2020, NR-52281. U.S. Department of Health and Human Services and Stanford ChEM-H/IMA. Details in supplementary text.

## Author Contributions

Study concept and design: AGB and VBM. Acquisition of data: YJS, GV, DP, KL, MO, SS. Data analysis and interpretation: YJS, GV, DP, KL, MO, SS, PBM, AGB, VBM. Drafting of the manuscript: YJS, GV, DP, PBM, AGB, VBM. Critical revision of the manuscript: PBM, AGB, VBM. Obtained funding: PBM, AGB, VBM. Administrative, technical, and material support: VBM. Study supervision: PBM, VBM, AGB. YJS, GV, DP, and KL contributed equally to this work. The authors declare no competing interests.

## Data and Materials Availability

Correspondence and requests for materials should be addressed to Vinit B. Mahajan. Reagents are available with a Materials Transfer Agreement. The raw docking data, parameters, and 3DPhyloFold code are deposited to Mendeley Data (DOI:10.17632/h3pmycddwc.1 and 10.17632/kk3gkzdsbf.2.)

## Methods

### Experimental model and subject details: mice, virus, and cells

Specific pathogen-free 6-week-old male and female BALB/c mice and were purchased from Envigo and maintained in the Animal Care Facilities at the University of Iowa. All protocols were approved by the Institutional Animal Care and Use Committees of the University of Iowa. The human serotype 5 adenoviral vector expressing human ACE2 under the control of the CMV promoter was previously described (VVC-McCray-7580; University of Iowa Viral Vector Core).^5^ The SARS-CoV-2 strains (SARS-Related Coronavirus 2 Isolate USA-WA1/2020) were obtained from BEI (Cat. # NR-52281) and Calu-3 2B4 cells (obtained from the Perlman Laboratory, University of Iowa). pVSV-ΔG-Luc was previously described.^18^ Calu-3 2B4 cells were grown in MEM (GIBCO, Grand Island, NY) supplemented with 20% FBS.

### Database search and sequence alignment

We first searched the UniProt database for reviewed entries denoted as transmembrane serine proteases (containing an S1-peptidase domain). This initial search yielded 9 manually curated sequences. A seed multiple sequence alignment (MSA) of S1-peptidase domains was then constructed using MAFFT v7 (alignment strategy: FFT-NS-1).^26^ Using HMMER-3.1 and the seed alignment, we produced an HMM profile and used it to broaden the search against the UniProt database (search restricted to reviewed sequences).^27^ This search yielded a total of 828 S1-peptidase sequences. We discarded fragmented sequences (<200 amino acids) that appeared too short to truly represent the S1-peptidase fold and redundant proteins were further filtered using CD-HIT v4 (100% threshold).^28^ This resulted in a pool of 742 proteins that were aligned using MAFFT v7 (alignment strategy FFT-NS-2).^26^ Sequences producing many gaps in the alignment were removed using MaxAlign, resulting in 600 S1-peptidase sequences.^29^

### Phylogenetic tree reconstruction

We used the IQ-TREE-1.6.2 algorithm to generate a maximum likelihood tree of the 600 S1-peptidase sequences.^30^ The IQ-TREE model finder tool was used to determine the best substitution model to fit the data. The Whelan & Goldman (WAG) substitution model was determined to be the best fit to the data. Bootstrap analysis was performed using the ‘ultra-fast’ method in IQ-TREE-1.6.2 with 1,000 replicas.

### Structural modeling of TMPRSS2-S1P

Briefly, a BLAST search of human TMPRSS2-S1P against the Protein Data Bank (PDB) returned the structure of human Hepsin (PDB 1Z8G) as the top hit. Other close matches were KLKB1 (PDB 6ESO), Plasminogen (PDB 4DUR), and Prostatin (PDB 3E16). A TMPRSS2-S1P model was generated with the Hepsin template (41% sequence identity) using Phyre2, MODELLER, and SWISS-Model. The models were in agreement and aligned well with minor variations in surface-exposed loop regions. The TMPRSS2-S1P model was then analyzed by ConSurf as previously described.^31^ The 600 sequences from our sequence-based phylogenetic analysis underwent MSA using MAFFT and conservation scores were calculated using the Bayesian method option in ConSurf. The TMPRSS2-S1P binding pocket was inferred by comparison to the structure of Hepsin bound to a peptidomimetic inhibitor (PDB 1Z8G) in PyMOL (The PyMOL Molecular Graphics System, Version 1.8 Schrödinger, LLC.).

### Structure-based phylogenetic analysis

There are over 2,000 structures of S1-peptidase domains represented in the PDB. We therefore searched the Pfam database for structures of mammalian peptidases and selected 74 representative structures (representing the wild-type protein) with an atomic resolution 3.2 Å or better (**Supplementary Table 2**).^32^ One structure per unique protein, fitting the above criteria, was selected. Structures (with reflection data deposited in the PDB) were evaluated by their reported global validation metrics in PDB-REDO.^33^ Re-refined structural models were used for further analysis. Structures were superimposed using PyMOL to calculate the pairwise root mean square deviation (RMSD) between protein alpha carbon atoms (Cα). A structural dissimilarity matrix (SDM) was constructed using the Cα RMSD values in order to generate a phylogenetic tree as previously described.^31^ To expedite the pairwise alignment process, we developed a Python-based script (named 3DPhyloFold) to perform the pairwise alignment of protein structures and generate an SDM. The phylogenetic tree was constructed using the UPGMA (Unweighted Pair Group Method with Arithmetic Mean) method in MEGAX software as previously described.^31^ For comparison, the sequences from the corresponding structures were also analyzed by sequence-based phylogeny. The 75 S1-peptidase sequences were aligned with MAFFT v7 ^26^ and analyzed in IQ-TREE-1.6.2.^30^ The Jones-Taylor-Thornton (JTT) substitution model was determined to be the best fit to the data. Bootstrap analysis was performed in IQ-TREE-1.6.2 (1,000 replicas). A TMPRSS2-S1P structure similarity score for each analyzed protease was calculated by dividing the pairwise sequence identity (to TMPRSS2-S1P) by the Cα RMSD of the pairwise alignment (**Extended Data Fig. 2**).

### Database search for S1-peptidase inhibitors

We first searched for inhibitors designed for the related proteins. We filtered for cases where a strong Structure-Activity-Relationship (SAR) between the ligand and protein was studied, where applicable. We discarded studies where the inhibitors displayed low potency, focusing on groups of inhibitors that displayed sub-micromolar inhibition for their intended protein. We focused on cases where the inhibitors studied contained a guanidine, or guanidine-like, functional group to interact with the S1 specificity pocket. Inhibitors were then prepared and docked against our TMPRSS2-S1P model.

### In silico docking calculations

Published crystal structures of inhibitor-bound Trypsin-3 (PDB 1H4W), KLKB1 (PDB 6O1S), and Factor VII (PDB 1W7X) were loaded into Maestro software (Schrödinger Release 2019-3). The TMPRSS2-S1P model described above was used. The protein preparation wizard was used to prepare the proteins for docking and simulations. The default parameters were used for the optimization of hydrogen-bond assignment (sampling of water orientations and use of pH 7.0). Waters molecules beyond 3 Å of heteroatoms or with fewer than three hydrogen bonds to non-waters were removed. Restrained energy minimization was applied using the OPLS3e force field. Prepared protein systems were further checked by Ramachandran plots, ensuring there were no steric clashes. To generate receptor grids for small molecule docking, the co-crystalized ligand was selected as the grid-defining ligand for each system. Default van der Waals radius scaling parameters were used (scaling factor of 1, partial charge cutoff of 0.25). For peptides, the grid size was made suitable for peptides to be docked. Default van der Waals radius scaling parameters were used (scaling factor of 1, partial charge cutoff of 0.25). For docking of the ligands into the various prepared proteins, the 3D structure was loaded into Maestro. Ligprep was used to prepare the ligands (by generating possible states at pH 7.0 ± 2.0 and retaining the specified stereochemical properties). The prepared small molecule ligands and peptide fragments were then docked using the most stringent docking mode (extra precision, “XP”) of Glide. Parameters and output files for the Glide runs can be found in Mendeley Data under the dataset identifier (DOI): 10.17632/h3pmycddwc.1.

### Docking Soybean Trypsin Inhibitor to TMPRSS2

The HADDOCK 2.4 online docking tool was used to generate TMPRSS2-S1P/SBTI complex structure model^34^. The TMPRSS2-S1P homology model and the SBTI structure (PDB 1AVW) were used for docking. To define the potential interaction surface between TMPRSS2 and SBTI, the TMPRSS2-S1P homology model was superimposed to the wild boar trypsin structure in complex with SBTI (PDB 1AVW) using PyMOL. The following residues of SBTI were designated as active residues: 501-502, 510, 512-514, 560-572, and 616-617. The overall Cα RMSD between the two models was 0.54 Å. SBTI was also docked to porcine Trypsin (PDB 1AVW), human Factor VII (PDB 1W7X), and human KLKB1 (PDB 6O1S). The HADDOCK scores represent the average score of the best cluster. The parameters and output files for the HADDOCK run can be found in Mendeley Data under the dataset identifier (DOI): 10.17632/h3pmycddwc.1.

### Protease activity array

Avoralstat, SBTI, PCI-27483, and Antipain were assessed for inhibition against TMPRSS2 and a panel of recombinant proteases by commercial services from Reaction Biology Corp. The Reaction Biology Corp profile tested in a 10-dose IC_50_ with, in triplicate, a 4-fold serial dilution starting at 10 μM against 11 proteases in **Fig. 3e** and a 3-fold serial dilution starting at 10 μM against 70 proteases in **Fig. 3f**. Compounds exhibit no fluorescent background that could interfere with the assay. The protease activities were monitored as a time-course measurement of the increase in fluorescence signal from fluorescently labeled peptide substrate, and initial linear portion of slope (signal/min) was analyzed.

### TMPRSS2-S1P expression and purification

The human TMPRSS2-S1P sequence (residues 252 - 489) was cloned into a pET28a vector with a N-terminal 6x-His tag. Plasmids were amplified and isolated from DH5α cells and transformed into *E. coli* BL21 (DE3). BL21 cells expressing TMPRSS2-S1P were induced with 0.5 mM IPTG. Cell pellets were resuspended in 35 to 50 mL of lysis buffer (50 mM Tris, 150 mM NaCl, 20 mM Imidazole pH 8.0, one tablet of EDTA-free protease inhibitor [Roche; Product # COEDTAF-RO], DNaseI [Roche; Product #11284932001]) and lysed and centrifuged for 30 minutes at 18,000 x g at 4 °C. Pellets were denaturated (50 mM Tris, 150 mM NaCl, 6 M Guanidinium Chloride, 1 M L-Arginine, 2 mM DTT pH 8.0), resuspended, and filtered with 0.22 μm filter. Refolding buffer-1 (50 mM Tris, 150 mM NaCl, 2 M Guanidinium Chloride, 1 M L-Arginine pH 8.0) was applied to SnakeSkin Dialysis Tubing (10,000 MWCO; Thermo Scientific™) and underwent refolding by dialyzing in 2 L of refolding buffer-1 at 4 °C. After the over-night refolding, the sample was filtered with 0.22 μm filter to remove aggregates and went through another step of dialysis in 2 L of refolding buffer-2 (50 mM Tris, 150 mM NaCl, 250 mM L-Arginine pH 8.0) for 1.5 hours at room-temperature. Sample was concentrated with a 10 kDa NMWL spin concentrator and passed over a HiLoad® 16/600 Superdex® 200 pg (GE Healthcare, Cat. # 28-9893-35) size-exclusion (SEC) column connected to an ÄTKA™ pure fast protein liquid chromatography (FPLC) system (GE Healthcare Inc.). The column was equilibrated with SEC buffer (50 mM Tris, 150 mM NaCl, pH 8.0). The final purity of recombinant TMPRSS2-S1P used for *in vitro* assays were >95% (**Extended Data Fig. 5a**).

### Measurement of TMPRSS2 activity

TMPRSS2-S1P proteolytic activity was confirmed by hydrolysis of the synthetic urokinase substrate, Cbz-GGR-AMC (Echelon Biosciences; Product #869-25). An enzyme titration in the presence of 50 μM Cbz-GGR-AMC revealed that maximal TMPRSS2-S1P activity occurred at high nanomolar (250 – 500 nM) protein concentrations (data not shown). The remaining assays were performed as followed: Briefly, 250 nM of purified TMPRSS2-S1P was added to a reaction buffer containing 50 mM Tris-HCl (pH 8.0), and 150 mM NaCl in black-bottom 96-well plates (100 μL per reaction). Inhibition experiments were carried out in the presence of 50 μM Cbz-GGR-AMC in the presence 10 to 500 μM compound: Camostat (Sigma-Aldrich; Cat. # SML0057), Avoralstat (MedChemExpress; Cat. # HY-16735), PCI-27483 (Cayman Chemical; Item #21334), Antipain (Sigma-Aldrich; Product #A6191), Leupeptin (Sigma-Aldrich; Product #L2884), MDL-28170 (Sigma-Aldrich; Product #M6690), Ritonavir (Sigma-Aldrich; Product #SML0491), or 5% DMSO (as a negative control). DMSO caused SBTI (Roche; Product #10109886001) to precipitate out of solution (unpublished observation). Inhibition experiments with SBTI (2 to 150 μM) were therefore performed in the absence of DMSO. Reactions were run at 37 °C for 30 minutes on a fluorimetric plate reader (Tecan Spark, Männedorf Switzerland). Proteolytic activity was measured as change in raw fluorescence units (ΔRFU; λexc = 373 nm, λem = 455 nm) at 30-second intervals. All experiments were performed in triplicate. The initial velocity (RFU/sec) of the reaction was measured by calculating the slope of the fluorescence data from the first three minutes. Kinetic parameters were then calculated by direct fitting to the Michaelis-Menten or Hill equation in GraphPad Prism 8 (GraphPad, San Diego, CA). There was no activity as expected with the cysteine protease substrate sLY-AMC (Bachem; Product #4002047; negative control; **Extended Data Fig. 5**).

### TMPRSS2 autoproteolysis assay

HEK 293T cells (ATCC^®^ Cat. # CRL-3216) were obtained from the Viral Vector Core Facility at the University of Iowa. Cells were grown in Dulbecco modified Eagle medium (DMEM) supplemented with 5% fetal bovine serum (Gibco), penicillin and streptomycin (Gibco, WT15140-122) and were maintained in a humidified atmosphere of 5% CO_2_ at 37 °C. Plasmid pEGFPN1 was obtained from Clontech. TMPRSS2-FL cDNA (pcDNA3.1-SARS-2-S-C9; obtained from the Gallagher Laboratory, Loyola University Medical Center, Illinois). Briefly, TMPRSS2-FL cDNA, containing a C-terminal anti-FLAG epitope tag, were amplified with PCR using pCMV-Sport6-TMPRSS2 template. The amplificates were cloned into pCAGGS.MCS via SacI and XhoI sites. The enzymatically-inactive pCAGGS-TMPRSS2(S441A)FLAG mutant cDNA was generated using QuickChange Site-Directed Mutagenesis Kit per manufacturer instructions (Agilent Technologies). Transient transfections of HEK-293T cells were performed using PolyFect transfection reagent per manufacturer instructions (Qiagen). For transfection, 2 μg of each plasmid (GFP [served as negative control], TMPRSS2 WT and S411A mutant) were dissolved in serum free media. PolyFect (20 μL) was added to the DNA solution followed by 10-minute incubation at room temperature. Growth media (0.6 mL) was then added to the reaction tubes and the transfection mix was immediately added onto the cells. 24 hours post-transfection, cell lysates were prepared using HNB buffer containing 0.1% protease inhibitor (Sigma-Aldrich, #P2714), incubated on ice for 20 minutes and centrifuged at 2,000 x g for 10 minutes. Supernatants were collected and protein concentration determined by DC protein assay reagent kit (BioRad). After separation by SDS-PAGE (4 to 12% Bis-Tris gradient gel), proteins were transferred to a PVDF membrane and blocked for 1-hr at room temperature using 5% nonfat dry milk in TBST. Membranes were probed with mouse monoclonal anti-Flag antibody (1:1,000; Sigma-Aldrich; Cat. #F3165) for 16 hours at 4° C. Blots were then washed three times with TBST (10 minutes/wash) and subsequently incubated with immunoglobulin-G labelled with horseradish peroxidase conjugated secondary anti-mouse antibody (1:5,000; Thermo Scientific™; Cat. #31432). Proteins were visualized by SuperSignal™ West Pico PLUS chemiluminescence reagent on a MyECL imager (Thermo Scientific™). Membranes were re-probed with β-actin (1:5000; Sigma-Aldrich, Cat. #A2228) as a loading control. The TMPRSS2-FL band intensity of each lane was normalized using the band intensity of corresponding β-actin loading control. Then, the normalized intensity of each lane was converted to the relative band intensity by comparison to the normalized band intensity of TMPRSS2-S441A in the same gel. The analyzed densitometry data from the total of 5 gel runs in **Fig. 3A** and **Extended Data Fig. 6** were combined. Data were analyzed by 1-way ANOVA followed by Dunnett’s multiple comparisons test using GraphPad Prism 8.0. Differences of p<0.0332 were considered statistically significant.

### Pseudovirus transduction assay

HEK-293T cells were transfected to express either the SARS-CoV-2 spike protein (with the cytoplasmic tail removed; residues 1 - 1255) or the full-length Vesicular Stomatitis Virus (VSV)-G protein. Then, these cells were transduced with a VSV vector expressing luciferase (VSV-ΔG-Luc), and pseudotyped with SARS-CoV-2 spike protein or VSV-G. After 2 hours at 37° C, the cells were washed 3 times to remove residual virus. Supernatant containing pseudovirus was harvested 3 times at 24-hour intervals and centrifuged to remove cellular debris. Pseudovirus from the 3 collections was pooled and ultracentrifuged through a 20% sucrose cushion for purification and concentration (100x). For the transduction assays, Calu-3 2B4 cells were grown in 96-well plates until confluent. Cells were incubated with the respective compounds for 1 hour at 37° C. After 1 hour, cells were transduced with pseudovirus, maintaining the same concentration of compounds, and incubated overnight. Transduction efficiency was assessed by quantifying luciferase activity in cell lysates using a commercial kit (Luciferase Assay System, Promega, Cat. #E1500) and a plate-reading luminometer (SpectraMax i3x, Molecular Devices). Data were analyzed by 2-way ANOVA followed by Dunnett’s multiple comparisons test using GraphPad Prism 8.0. Differences of p<0.0332 were considered statistically significant.

### Infectious SARS-CoV-2 neutralization assay

The 2019n-CoV/USA-WA1/2019 strain of SARS-CoV-2 (Accession number: MT985325.1) used in these studies was passaged on Calu-3 2B4 cells and sequence verified. Calu-3 2B4 cells were plated in 48 well plates. Cells were incubated with medium containing indicated compounds or vehicle for 1 hour at 37° C. The medium was removed and SARS-CoV-2 (MOI=0.1) in medium containing indicated compounds were added into each well. The cells were incubated with viruses for 1 hour at 37 °C. Next, the viruses were removed, and cells were rinsed with PBS once to remove remaining viruses. After that, cells were incubated with medium containing indicating compounds overnight. Following day, Total cellular RNA was isolated using Directzol RNA MiniPrep kit (Zymo Research, Cat. # R2052) from TRIzol (Invitrogen; Cat. #15596018). A DNase treatment step was included. Total RNA (500 ng) was used for cDNA syntheses by High-Capacity cDNA Reverse Transcription Kit (Applied Biosystems; Cat. # 4368814). Realtime PCR was applied to quantify viral genomic RNA and HPRT mRNA levels (SARS-2-N1-F primer: GACCCCAAAATCAGCGAAAT; SARS-2-N1-R primer: TCTGGTTACTGCCAGTTGAATCTG; Human HPRT-F primer: AGGATTTGGAAAGGGTGTTTATTC; Human HPRT-R primer: CAGAGGGCTACAATGTGATGG; Integrated DNA Technologies). The relative abundance of viral genomic RNA normalized to HPRT was calculated and presented as 2^-ΔCT^. All the data were analyzed using GraphPad Prism 8.0. Data were analyzed by 2-way ANOVA followed by Dunnett’s multiple comparisons test. Differences of p < 0.0332 were considered statistically significant.

### Transduction and infection of mice

Mice were anesthetized with ketamine/xylazine (87.5 mg/kg ketamine/12.5 mg/kg xylazine) and transduced intranasally with 2.5 x 10^8^ FFU of Ad5-ACE2 in 75 mL DMEM. Five days post transduction, mice were infected intranasally with SARS-CoV-2 (3 x 10^3^ or 1 x 10^5^ PFU) in a total volume of 50 mL DMEM. Infected mice were treated with Avoralstat, Camostat (30 mg/kg intraperitoneal injection), or vehicle (DMSO; negative control) either four hours before and after being challenged by virus, or two doses per day (8 to 9 hours apart) for three days post infection. Virus titers were measured in harvested lungs by plaque assay 1-day post infection. The weight was monitored for 6 days post infection. Data were analyzed by 2-way ANOVA followed by Dunnett’s multiple comparisons test. Differences of p<0.0332 were considered statistically significant.

### SARS-CoV-2 plaque assay

Lung homogenate supernatants were serially diluted in DMEM. Vero E6 cells in 12 well plates were inoculated at 37 ºC in 5% CO_2_ for 1 hour with gentle rocking every 15 minutes. After removing the inocula, plates were overlaid with 1.2% agarose containing 10% FBS. After further incubation for 3 days, overlays were removed, and plaques were visualized by staining with 0.1% crystal violet. Viral titers were calculated as plaque forming units (PFU) per lung. All work with SARS-CoV-2 was conducted in the Biosafety Level 3 (BSL3) Laboratories of the University of Iowa. These studies were approved by the University of Iowa Institutional Animal Care and Use Committee.

## References

1 Hoffmann, M. et al. SARS-CoV-2 Cell Entry Depends on ACE2 and TMPRSS2 and Is Blocked by a Clinically Proven Protease Inhibitor. Cell, doi:10.1016/j.cell.2020.02.052 (2020).

2 Kawase, M., Shirato, K., van der Hoek, L., Taguchi, F. & Matsuyama, S. Simultaneous treatment of human bronchial epithelial cells with serine and cysteine protease inhibitors prevents severe acute respiratory syndrome coronavirus entry. J Virol 86, 6537–6545, doi:10.1128/JVI.00094-12 (2012).

3 Iwata-Yoshikawa, N. et al. TMPRSS2 Contributes to Virus Spread and Immunopathology in the Airways of Murine Models after Coronavirus Infection. J Virol 93, doi:10.1128/JVI.01815-18 (2019).

4 Shrimp, J. H. et al. An Enzymatic TMPRSS2 Assay for Assessment of Clinical Candidates and Discovery of Inhibitors as Potential Treatment of COVID-19. ACS Pharmacol Transl Sci 3, 997–1007, doi:10.1021/acsptsci.0c00106 (2020).

5 Sun, J. et al. Generation of a Broadly Useful Model for COVID-19 Pathogenesis, Vaccination, and Treatment. Cell, doi:10.1016/j.cell.2020.06.010. (2020).

6 Riedl, M. A. et al. Evaluation of avoralstat, an oral kallikrein inhibitor, in a Phase 3 hereditary angioedema prophylaxis trial: The OPuS-2 study. Allergy 73, 1871–1880, doi:10.1111/all.13466 (2018).

7 Zhou, P. et al. A pneumonia outbreak associated with a new coronavirus of probable bat origin. Nature 579, 270–273, doi:10.1038/s41586-020-2012-7 (2020).

8 Kim, T. S., Heinlein, C., Hackman, R. C. & Nelson, P. S. Phenotypic analysis of mice lacking the Tmprss2-encoded protease. Mol Cell Biol 26, 965–975, doi:10.1128/MCB.26.3.965-975.2006 (2006).

9 Shirato, K., Kawase, M. & Matsuyama, S. Middle East respiratory syndrome coronavirus infection mediated by the transmembrane serine protease TMPRSS2. J Virol 87, 12552–12561, doi:10.1128/JVI.01890-13 (2013).

10 Shen, L. W., Mao, H. J., Wu, Y. L., Tanaka, Y. & Zhang, W. TMPRSS2: A potential target for treatment of influenza virus and coronavirus infections. Biochimie 142, 1–10, doi:10.1016/j.biochi.2017.07.016 (2017).

11 Midgley, I. et al. Metabolic fate of 14C-camostat mesylate in man, rat and dog after intravenous administration. Xenobiotica 24, 79–92, doi:10.3109/00498259409043223 (1994).

12 Talevi, A. & Bellera, C. L. Challenges and opportunities with drug repurposing: finding strategies to find alternative uses of therapeutics. Expert Opin Drug Discov 15, 397–401, doi:10.1080/17460441.2020.1704729 (2020).

13 Schweinitz, A. et al. Design of novel and selective inhibitors of urokinase-type plasminogen activator with improved pharmacokinetic properties for use as antimetastatic agents. J Biol Chem 279, 33613–33622, doi:10.1074/jbc.M314151200 (2004).

14 Jin, Z. et al. Structure of M(pro) from SARS-CoV-2 and discovery of its inhibitors. Nature 582, 289–293, doi:10.1038/s41586-020-2223-y (2020).

15 Shin, D. et al. Papain-like protease regulates SARS-CoV-2 viral spread and innate immunity. Nature 587, 657–662, doi:10.1038/s41586-020-2601-5 (2020).

16 Johnson, B. A. et al. Furin Cleavage Site Is Key to SARS-CoV-2 Pathogenesis. bioRxiv, doi:10.1101/2020.08.26.268854 (2020).

17 Shulla, A. et al. A transmembrane serine protease is linked to the severe acute respiratory syndrome coronavirus receptor and activates virus entry. J Virol 85, 873–882, doi:10.1128/JVI.02062-10 (2011).

18 Whitt, M. A. Generation of VSV pseudotypes using recombinant DeltaG-VSV for studies on virus entry, identification of entry inhibitors, and immune responses to vaccines. J Virol Methods 169, 365–374, doi:10.1016/j.jviromet.2010.08.006 (2010).

19 Schneider, C. A. et al. [Assessing myocardial viability in chronic myocardial infarct with 18F-fluoro-D-glucose positron emission tomography and 99mTc-MIBI SPECT]. Z Kardiol 83, 124–131 (1994).

20 Choi, J. Y. et al. Nafamostat Mesilate as an Anticoagulant During Continuous Renal Replacement Therapy in Patients With High Bleeding Risk: A Randomized Clinical Trial. Medicine (Baltimore) 94, e2392, doi:10.1097/MD.0000000000002392 (2015).

21 Cornpropst, M. et al. Safety, pharmacokinetics, and pharmacodynamics of avoralstat, an oral plasma kallikrein inhibitor: phase 1 study. Allergy 71, 1676–1683, doi:10.1111/all.12930 (2016).

22 Estimating the Maximum Safe Starting Dose in Initial Clinical Trials for Therapeutics in Adult Healthy Volunteers <https://www.fda.gov/regulatory-information/search-fda-guidance-documents/estimating-maximum-safe-starting-dose-initial-clinical-trials-therapeutics-adult-healthy-volunteers> (2005).

23 Song, H. K. & Suh, S. W. Kunitz-type soybean trypsin inhibitor revisited: refined structure of its complex with porcine trypsin reveals an insight into the interaction between a homologous inhibitor from Erythrina caffra and tissue-type plasminogen activator. J Mol Biol 275, 347–363, doi:10.1006/jmbi.1997.1469 (1998).

24 Bakowski, M. A. et al. Oral drug repositioning candidates and synergistic remdesivir combinations for the prophylaxis and treatment of COVID-19. bioRxiv, 2020.2006.2016.153403, doi:10.1101/2020.06.16.153403 (2020).

25 Fragkou, P. C. et al. Review of trials currently testing treatment and prevention of COVID-19. Clin Microbiol Infect, doi:10.1016/j.cmi.2020.05.019 (2020).

26 Katoh, K. & Standley, D. M. MAFFT multiple sequence alignment software version 7: improvements in performance and usability. Mol Biol Evol 30, 772–780, doi:10.1093/molbev/mst010 (2013).

27 Finn, R. D., Clements, J. & Eddy, S. R. HMMER web server: interactive sequence similarity searching. Nucleic Acids Res 39, W29–37, doi:10.1093/nar/gkr367 (2011).

28 Li, W. & Godzik, A. Cd-hit: a fast program for clustering and comparing large sets of protein or nucleotide sequences. Bioinformatics 22, 1658–1659, doi:10.1093/bioinformatics/btl158 (2006).

29 Gouveia-Oliveira, R., Sackett, P. W. & Pedersen, A. G. MaxAlign: maximizing usable data in an alignment. BMC Bioinformatics 8, 312, doi:10.1186/1471-2105-8-312 (2007).

30 Nguyen, L. T., Schmidt, H. A., von Haeseler, A. & Minh, B. Q. IQ-TREE: a fast and effective stochastic algorithm for estimating maximum-likelihood phylogenies. Mol Biol Evol 32, 268–274, doi:10.1093/molbev/msu300 (2015).

31 Velez, G. et al. Structural Insights into the Unique Activation Mechanisms of a Non-classical Calpain and Its Disease-Causing Variants. Cell Rep 30, 881–892 e885, doi:10.1016/j.celrep.2019.12.077 (2020).

32 Finn, R. D. et al. Pfam: the protein families database. Nucleic Acids Res 42, D222–230, doi:10.1093/nar/gkt1223 (2014).

33 Joosten, R. P. et al. PDB_REDO: automated re-refinement of X-ray structure models in the PDB. J Appl Crystallogr 42, 376–384, doi:10.1107/S0021889809008784 (2009).

34 de Vries, S. J., van Dijk, M. & Bonvin, A. M. The HADDOCK web server for data-driven biomolecular docking. Nat Protoc 5, 883–897, doi:10.1038/nprot.2010.32 (2010).

