## Supplementary Information for "TMPRSS2 structure-phylogeny repositions Avoralstat for SARS-CoV-2 prophylaxis in mice"

### *Additional acknowledgements*

At Stanford University, we thank Jing Yang, Angela S. Li, and Teja Chemudupati in the Department of Ophthalmology for technical assistance; Dr. Jeffrey L. Goldberg (Chair of Ophthalmology) for supporting our research during the COVID-19 campus lockdown; and Dr. Corey Liu and Dr. Daniel Fernandez in the Stanford ChEM-H Macromolecular Structure Knowledge Center for supporting protein production along with Dr. Mark Smith (Director) in the Stanford ChEM-H Medicinal Chemistry Knowledge Center for helpful discussions. From the University of Iowa, we thank Shu Wu in the Department of Pediatrics for technical assistance; and we thank Dr. Stanley Perlman for helpful discussions. From Loyola University Chicago, we thank Dr. Thomas Gallagher for sharing TMPRSS2 plasmids.

### *Funding information*

VBM and AGB are supported by NIH grants [R01EY026682, R01EY024665, R01EY025225, R01EY024698, and P30EY026877], and Research to Prevent Blindness, New York, New York. GV is supported by NIH grants [F30EYE027986 and T32GM007337]. DP is supported by a NEI/NIH T32EY027816 training grant. The study was also supported by P01AI060699 (PBM); and the Pathology Core, which is partially supported by the Center for Gene Therapy for Cystic Fibrosis (NIH Grant P30DK-54759), and the Cystic Fibrosis Foundation. PBM is supported by the Roy J. Carver Charitable Trust. MO is supported by a NHLBI/NIH T32HL007638 training grant.

### *In silico data and materials availability*

3DPhyloFold is open source and available at Mendeley Data under the dataset identifier (DOI): 10.17632/kk3gkzdsbf.2. The implementation notes, code, and description of methodology are available on the site. The raw docking data and parameters have been deposited to Mendeley Data with the dataset identifier (DOI): 10.17632/h3pmycddwc.1.

### *Quantification and statistical analysis in each figure data*

**Fig. 3: (a)** Densitometry analysis. The measured band intensity data from the total of 5 gel runs in panel A and **Extended Data Fig. 6** were combined. (n=14 for vehicle and Camostat, respectively; n=5 for each group of GFP, Avoralstat, PCI-27483, Antipain, and SBTI). Data represent the mean ± SEM from two independent experiments analyzed by 1-way ANOVA followed by Dunnett's multiple comparisons test (*p<0.0332, **p<0.0021, ***p<0.0002). **(c)** Calu-3 cells were pre-incubated with the indicated concentrations of DMSO (vehicle; negative control), Camostat, Avoralstat, PCI-27483, SBTI, and Antipain and subsequently inoculated with pseudovirus particles harboring SARS-CoV-2 Spike protein. Data represent the mean ± SEM (n = 6; number of technical replicates) and are fit to the Hill equation (Camostat $R^2$ = 0.71; Avoralstat $R^2$ = 0.74; PCI-27483 $R^2$ = 0.20; SBTI $R^2$ = 0.49; Antipain $R^2$ = 0.48). **(d)** SARS-CoV-2 viral gRNA in the presence of 100 μM of DMSO (vehicle; negative control) or inhibitor. Data represent the mean ± SEM (n = 3; number of technical replicates) and were analyzed by 1-way ANOVA followed by Tukey's multiple comparisons test (****$p<0.0001$ compared to vehicle). **(e)** SARS-CoV-2 viral gRNA as a function of Camostat or Avoralstat concentration. Data represent the mean ± SEM (n = 3; number of technical replicates) were analyzed by 2-way ANOVA followed by Dunnett's multiple comparisons test (*$p<0.0332$, ****$p<0.0001$ compared to vehicle).

**Fig. 4: (a)** Wild-type BALB/c mice transduced with Ad5-hACE2 were intranasally infected with $3 \times 10^3$ PFU of SARS-CoV-2. Mice were treated with Avoralstat, Camostat (30 mg/kg intraperitoneal injection), or vehicle (DMSO; negative control) four hours before and after being challenged by virus. Virus titers were measured in harvested lungs 1-day post infection. Data are represented as mean ± SEM (n = 3; number of mice) and were analyzed by 1-way ANOVA followed by Tukey's multiple comparisons test (*$p<0.05$; **$p<0.01$). **(v)** Wild-type BALB/c mice transduced with Ad5-hACE2 were intranasally infected with $1 \times 10^5$ PFU of SARS-CoV-2. Mice were treated with Avoralstat, Camostat (30 mg/kg intraperitoneal injection), or vehicle (DMSO; negative control) four hours before and after being challenged by virus. Virus titers were measured in harvested lungs 1-day post infection. Data are represented as mean ± SEM (n = 3; number of mice) and were analyzed by 1-way ANOVA followed by Tukey's multiple comparisons test (*$p<0.05$; **$p<0.01$). **(c)** Virus titers were measured in harvested lungs 5 d.p.i.. Data are represented as mean ± SEM and were analyzed by 1-way ANOVA followed by Tukey's multiple comparisons test (*$p<0.0332$; **$p<0.0021$, ***$p<0.0002$, ****$p<0.0001$ compared to vehicle; n = 4 for each group). **(d, f)** Weight change of the infected Ad5-hACE2 transduced mice for 10 days following treatment with Avoralstat, Camostat, or vehicle. Data are represented as mean ± SEM (n = 6; number of mice) and were analyzed by 2-way ANOVA followed by Dunnett's multiple comparisons test (*$p<0.0332$, ***$p<0.0002$ compared to vehicle). **(e)** Virus titers were measured in harvested lungs 2 and 4 d.p.i.. Data are represented as mean ± SEM and were analyzed by 1-way ANOVA followed by Tukey's multiple comparisons test (*$p<0.0332$; **$p<0.0021$, ***$p<0.0002$, ****$p<0.0001$ compared to vehicle; n = 3 for each group).

**Extended Data Fig. 5: (c)** Michaelis-Menten analysis of Cbz-GGR-AMC hydrolysis by 250 nM TMPRSS2-S1P in the presence of 5% DMSO. Data is displayed as mean ± SEM (n = 3; number of reactions) and fit to the Michaelis-Menten equation ($R^2 = 0.88$).

**Extended Data Fig. 7: (a)** SARS-CoV-2 and MERS-CoV viral gRNA as a function of Camostat concentration. Data represent the mean ± SEM (n = 3; number of technical replicates) were analyzed by 2-way ANOVA followed by Dunnett's multiple comparisons test (*$p<0.0332$, ****$p<0.0001$ compared to vehicle). **(b)** SARS-CoV-2 and MERS-CoV viral gRNA as a function of Avoralstat concentration. Data represent the mean ± SEM (n = 3; number of technical replicates) were analyzed by 2-way ANOVA followed by Dunnett's multiple comparisons test (*$p<0.0332$, **$p<0.0021$, ****$p<0.0001$ compared to vehicle).
