## Extended Data Figures for "TMPRSS2 structure-phylogeny repositions Avoralstat for SARS-CoV-2 prophylaxis in mice"

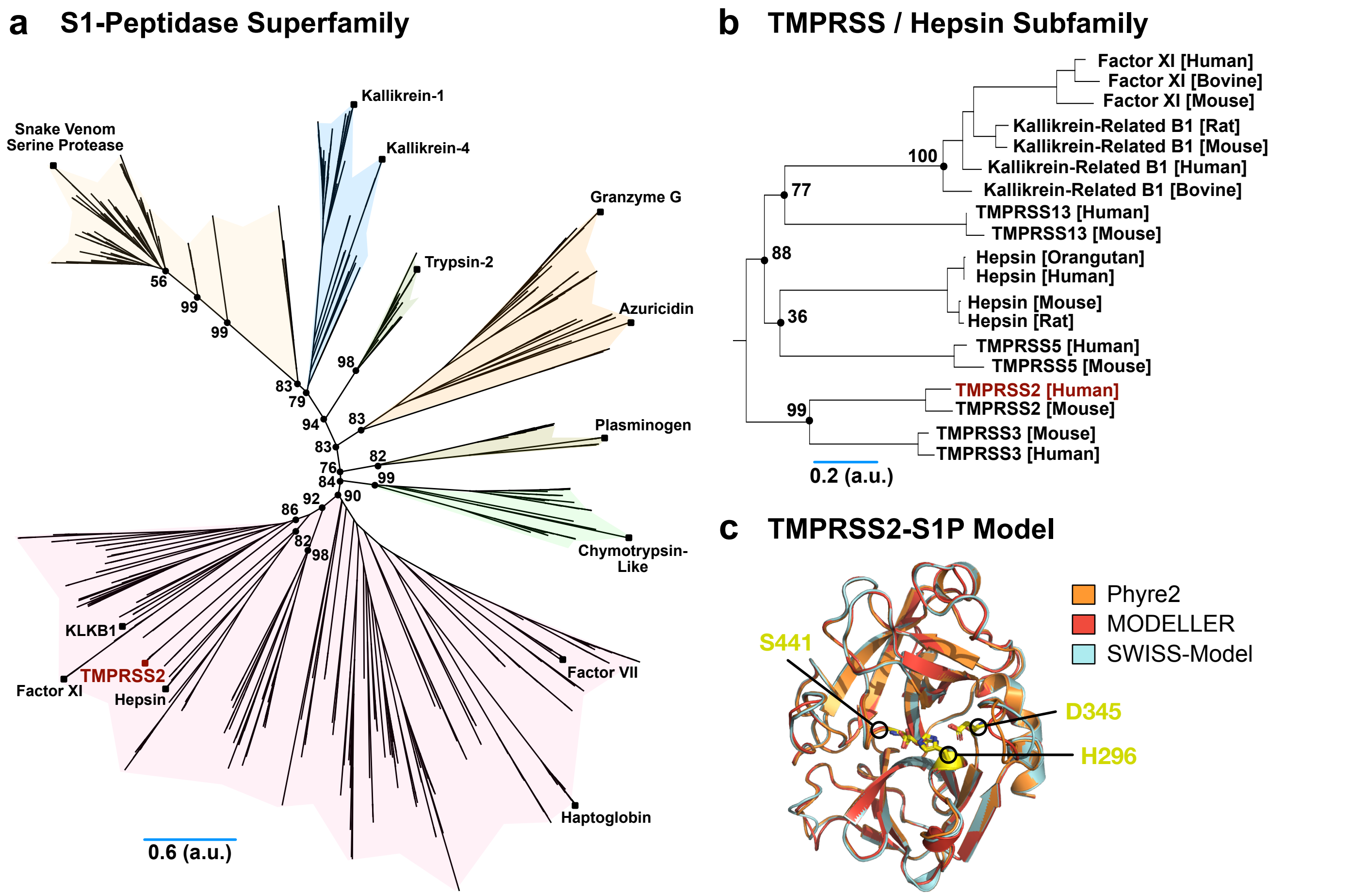


**Extended Data Fig. 1: Sequence-based phylogenetic analysis of the S1-peptidase superfamily reveals the evolutionary relatedness of TMPRSS2-S1P to other proteases**

We aligned a seed group of manually curated sequences from transmembrane serine proteases that share the S1-peptidase fold and produced a profile hidden Markov model (HMM). The profile HMM was then used in a HMMER search against the UniProt database to broaden the pool of S1-peptidase sequences to six hundred, which were then aligned. The enriched multiple sequence alignment (MSA) subsequently underwent rigorous phylogenetic reconstruction in IQ-TREE-1.6.2. **(a)** The reconstructed S1-peptidases superfamily tree representing 600 sequences across all species is shown. Evolutionary distances were inferred using the maximum likelihood method with 1,000 bootstrap replicas. The unrooted tree is represented with the main clades highlighted in different colors. Bootstrap values are projected onto the tree topology. The positions of TMPRSS2-S1P and representative serine proteases are denoted by black squares in each clade for reference. **(b)** The TMPRSS2-S1P/Hepsin subtree (bootstrap 82%; 19 sequences). Bootstrap values are projected onto the tree topology. **(c)** Ribbon tracing diagram representing the S1-peptidase domain of human TMPRSS2-S1P. The homology model was generated with three separate modeling programs (MODELLER, SWISS-Model, and Phyre2) using the structure of human Hepsin (PDB: 1Z8G) as a template. The catalytic triad residues (H296, D345, and S441) are represented by the yellow stick model. See also **Supplementary Table 1**.

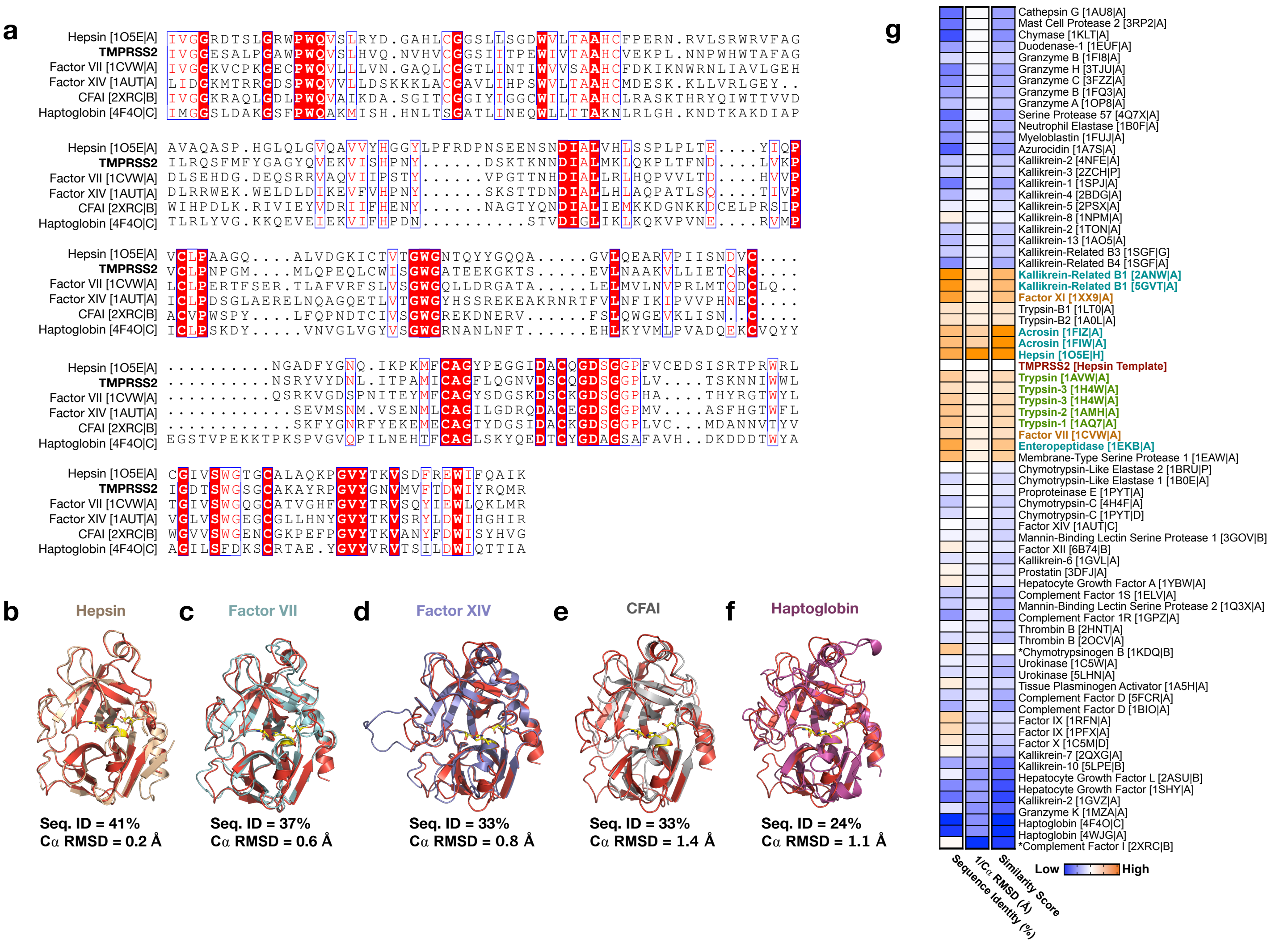


**Extended Data Fig. 2. Comparison of S1-peptidase structures**

**(a)** Multiple sequence alignment of representative peptidase domains from the structural phylogenetic analysis. The sequence alignment was generated in MAFFT and visualized using ESpript. **(b)** Overlay between Hepsin (PDB 1O5E), **(c)** Factor VII (PDB 1CVW), **(d)** Factor XIV (1AUT), **(e)** CFAI (2XRC), **(f)** Haptoglobin (4F4O) and TMPRSS2 (cyan). The corresponding sequence identity and Cα RMSD is represented below the structural alignment. **(g)** A heatmap representing the sequence and structural similarity to TMPRSS2-S1P is displayed to the right of the tree. The first column denotes the pairwise sequence identity (%) to TMPRSS2-S1P and the second column denotes the structural similarity (1/Cα RMSD) to the TMPRSS2-S1P model. The last column denotes the structural similarity 'score' which is calculated by dividing the pairwise sequence identity by the backbone RMSD.

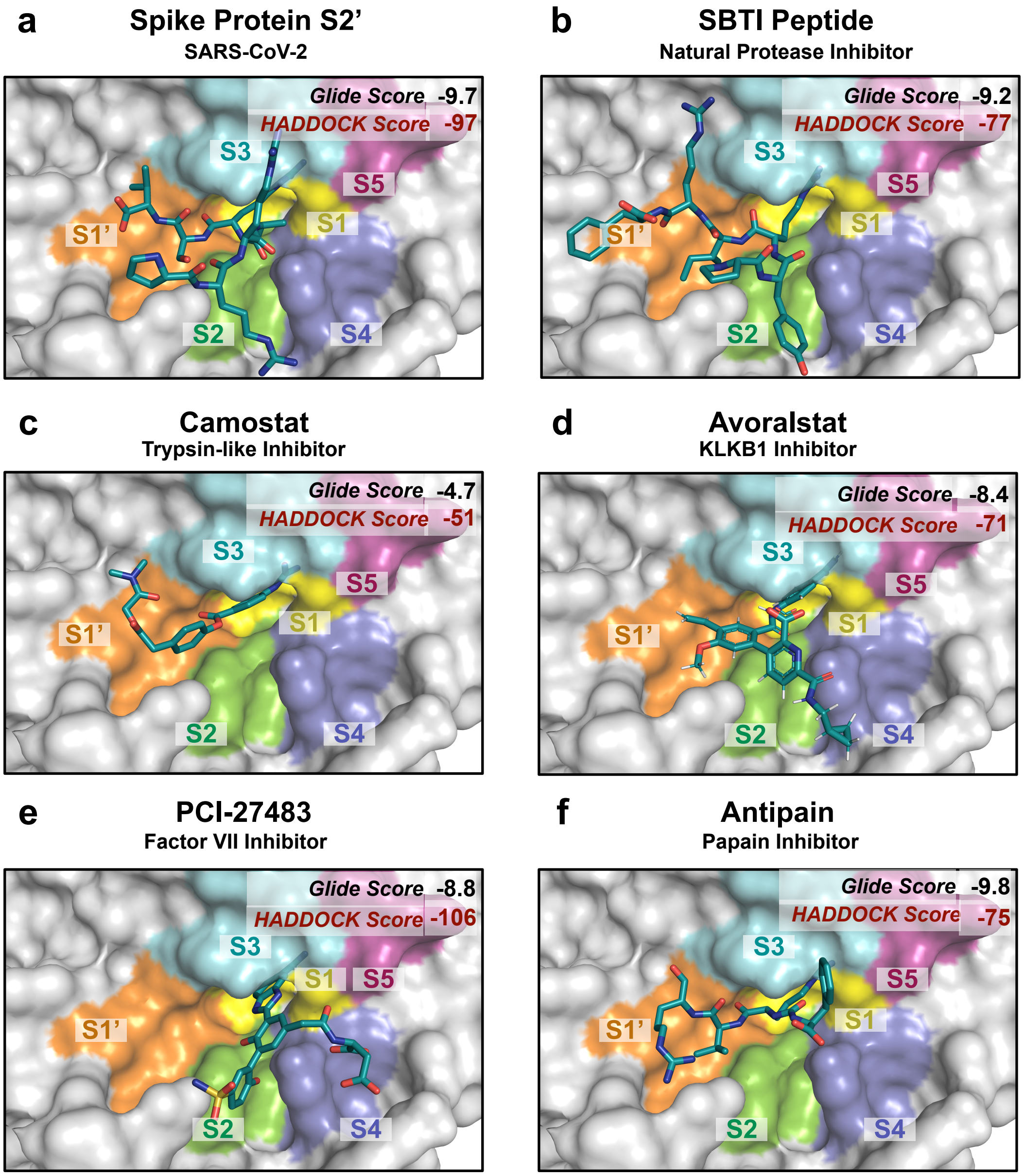


**Extended Data Fig. 3. Binding poses of the top scoring inhibitors**

**(a)** Spike S2’ peptide motif, **(b)** SBTI peptide motif, **(c)** Camostat, **(d)** Avoralstat, **(e)** PCI-27483, and **(f)** Antipain all binding strongly within the S1 sup-pocket of TMPRSS2. Each color represents different inhibitor binding sub-pockets: Orange; S1’ sub-pocket, yellow; S1 sub-pocket, green; S2 sub-pocket, cyan; S3 sub-pocket, slate; S4 sub-pocket, and violet; S5 sub-pocket. See also **Supplementary Table 3 and 4**.

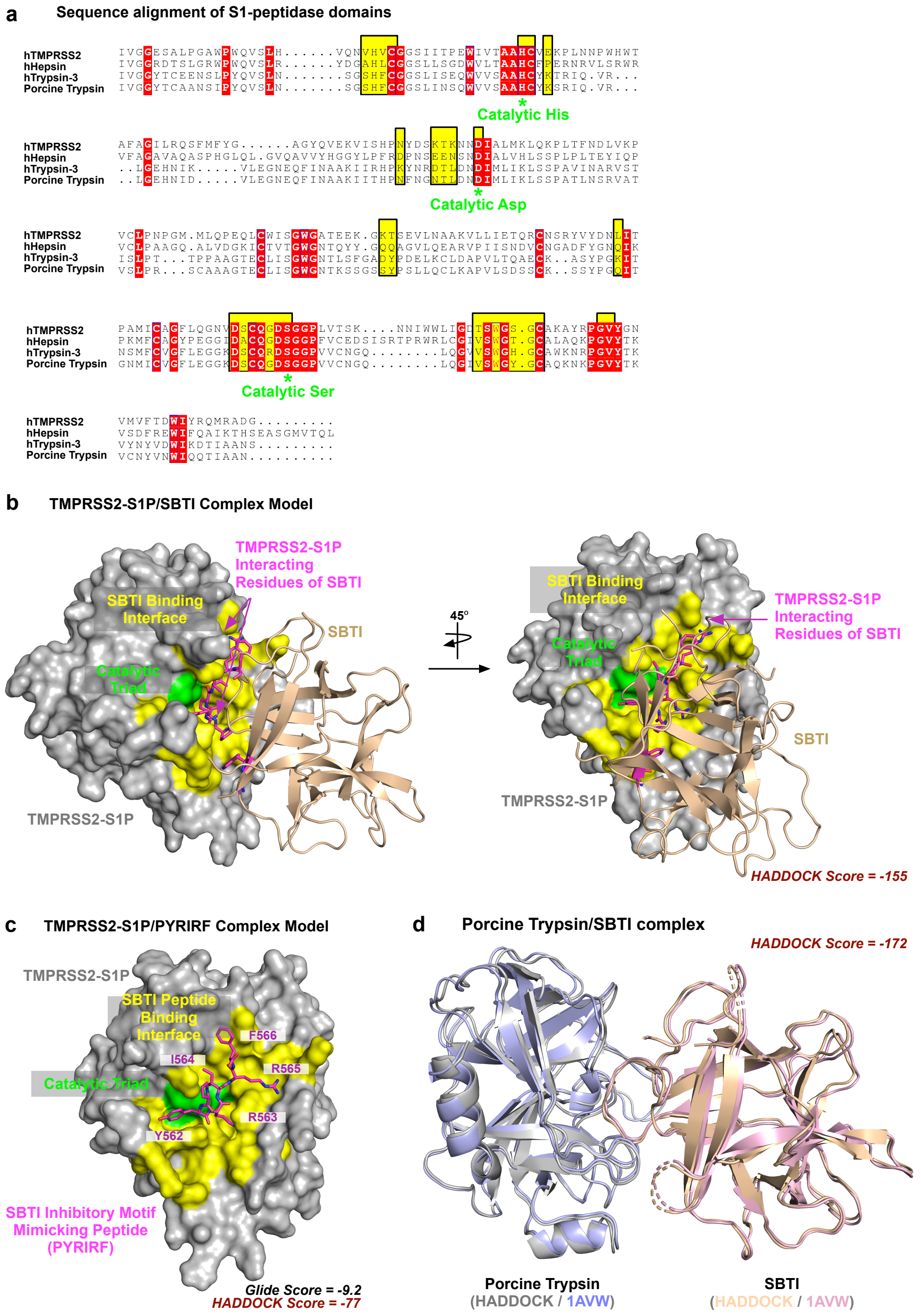


**Extended Data Fig.4. Soybean Trypsin Inhibitor (SBTI) derivatives might inhibit TMPRSS2-S1P activity** **(a)** Multiple sequence alignment of serine peptidase domains that were used for the binding pocket analysis. The sequence alignment was generated in MAFFT and visualized using ESPript 3.0. Residues involved in the SBTI binding interface are highlighted in yellow. * *- indicates the residues forming catalytic triad.* **(b)** Model of TMPRSS2-S1P in complex with SBTI generated by HADDOCK. Presented structure is one of the best structures in the biggest size and the best Z-score cluster. **(c)** SBTI Residues at the TMPRSS2-S1P/SBTI Interface are shown in the magenta stick model (residues 561-566; PYRIRF). Catalytic residues are shown in green, and R563 is the P1 residue. **(d)** Model of porcine Trypsin in complex with SBTI generated by HADDOCK.

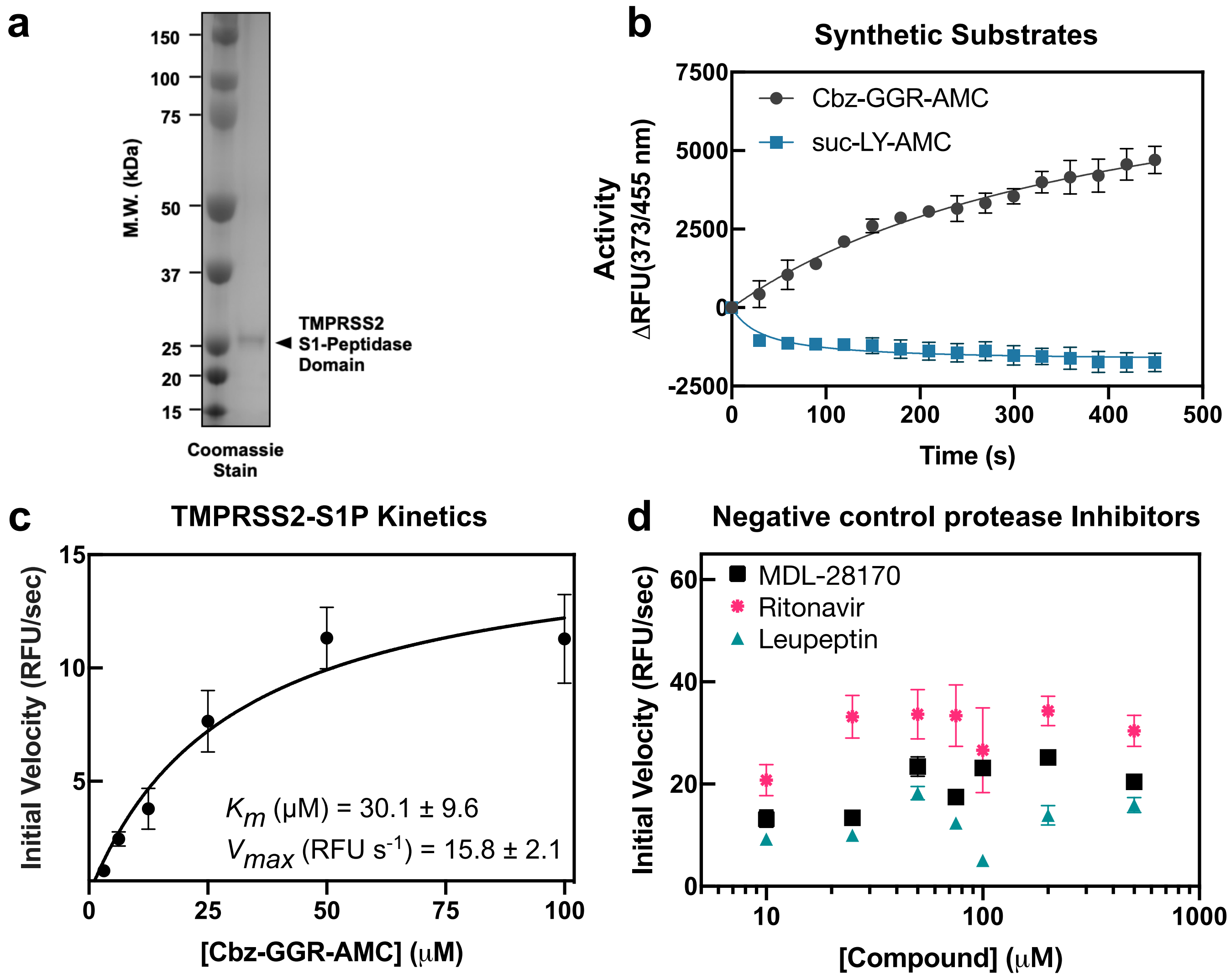


**Extended Data Fig. 5. Purification and *in vitro* characterization of TMPRSS2-S1P**

**(a)** Coomassie-stained SDS-PAGE gel of purified recombinant TMPRSS2-S1P. The protein was purified using affinity (nickel-NTA) and size-exclusion chromatography. **(b)** Fluorescence tracing of 50 μM Cbz-GGR-AMC (grey) or suc-LY-AMC (blue) in the presence of 250 nM TMPRSS2-S1P. Data represent the mean ± SEM of three technical replicates. **(c)** Michaelis-Menten analysis recombinant TMPRSS2-S1P hydrolysis of Cbz-GGR-AMC (fluorescence resonance energy transfer assay; data displayed as mean ± SEM; n=3). **(d)** Cbz-GGR-AMC (50 μM) hydrolysis by 250 nM TMPRSS2-S1P in the presence of 10-500 μM Leupeptin (black), MDL-28170 (magenta) and Ritonavir (blue/green). The initial velocity for each condition was plotted against the inhibitor concentration. Data represent the mean ± SEM of three technical replicates.

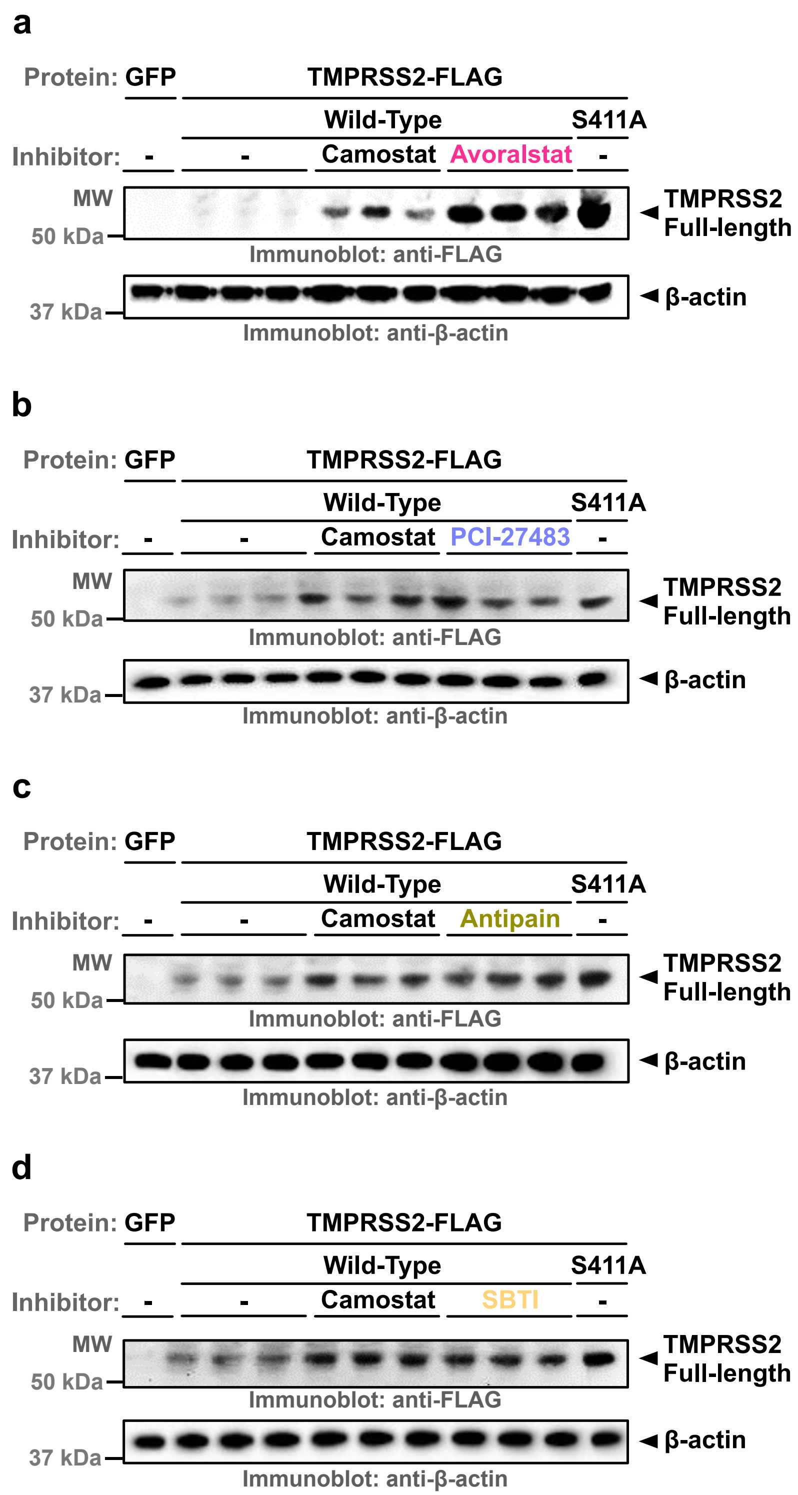


**Extended Data Fig. 6. Candidate molecules have similar efficacy as Camostat in inhibiting full length TMPRSS2 protease activity at 100 μM dose in cells**

Western-blot results of TMPRSS2-FL autoproteolysis assay by each compound compared with vehicle and Camostat: **(a)** Avoralstat; **(b)** PCI-27483; **(c)** Antipain; and **(d)** SBTI. Control vector (GFP) and TMPRSS2-S441A expressing vector were served as negative and positive controls, respectively.

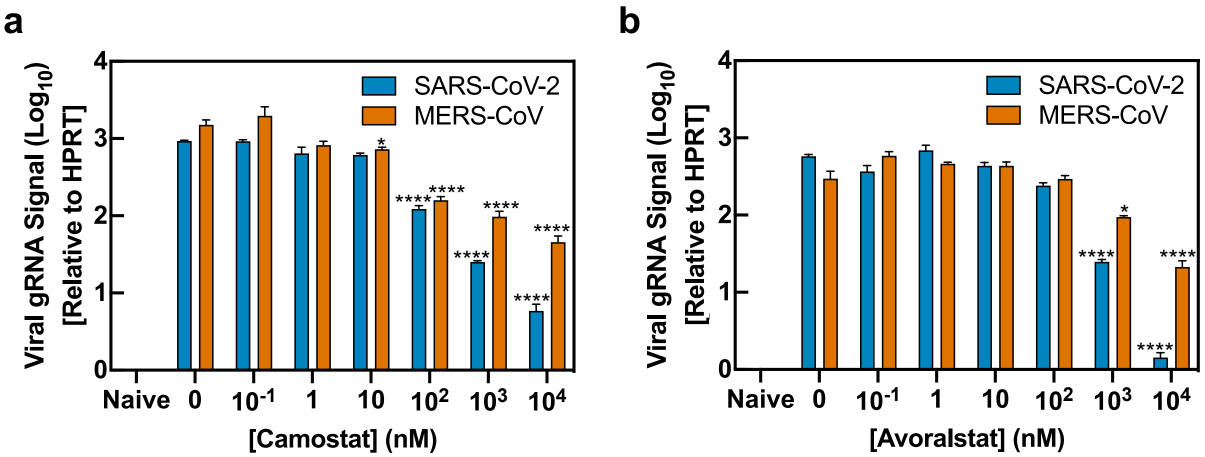


**Extended Data Fig. 7. SARS-CoV-2 is more sensitive to Avoralstat and Camostat than MERS-CoV**

Calu-3 cells were pre-incubated with the indicated concentrations of **(a)** Camostat or **(b)** Avoralstat and subsequently inoculated with SARS-CoV-2 or MERS-CoV (multiplicity of infection, MOI = 0.1) in inhibitor-containing media. Cells were subsequently washed, and viral gRNA was determined by qRT-PCR. Data represent the mean ± SEM were analyzed by 2-way ANOVA followed by Dunnett's multiple comparisons test (*$p<0.0332$, ****$p<0.0001$ compared to vehicle).
