## Supplementary Tables for "TMPRSS2 structure-phylogeny repositions Avoralstat for SARS-CoV-2 prophylaxis in mice"

| Position | Amino Acid | MSA Conservation (%) | TMPRSS2 Function |
|---|---|---|---|
| 259 | G | 99.3 | - |
| 270 | Q | 60.8 | - |
| 281 | C | 95.0 | S1’ sub-binding pocket |
| 293 | T | 81.2 | - |
| 294 | A | 95.0 | - |
| 295 | A | 92.8 | - |
| 296 | H | 92.3 | Catalytic His residue; Active site |
| 297 | C | 95.8 | S1’ sub-binding pocket |
| 313 | G | 97.5 | - |
| 345 | D | 99.0 | Catalytic Asp residue; Active site |
| 347 | A | 45.8 | - |
| 351 | L | 93.5 | - |
| 366 | L | 93.3 | - |
| 383 | G | 99.2 | - |
| 384 | W | 92.0 | - |
| 385 | G | 98.5 | - |
| 397 | L | 74.8 | - |
| 410 | C | 100 | - |
| 426 | C | 100 | - |
| 427 | A | 86.8 | - |
| 437 | C | 94.2 | S1 sub-binding pocket |
| 440 | D | 98.5 | S1’ sub-binding pocket |
| 441 | S | 93.3 | Catalytic Ser residue; Active site |
| 442 | G | 99.3 | - |
| 443 | G | 94.8 | - |
| 444 | P | 94.8 | - |
| 445 | L | 78.7 | - |
| 457 | G | 99.3 | - |
| 460 | S | 92.0 | S2 sub-binding pocket |
| 465 | C | 99.7 | S1 sub-binding pocket |
| 471 | P | 84.5 | - |
| 474 | Y | 85.3 | S1 sub-binding pocket |
| 483 | W | 100 | - |
| 484 | I | 88.2 | - |

**Supplementary Table 1. Conserved residues in S1-peptidases**

Conserved residues in 600 S1-peptidases across all species. Amino acids are numbered according their position in human TMPRSS2-S1P.

| Protein | Species | Residues | PDB | Chain | PDB residues | Resolution (Å) | Ligand? |
|---|---|---|---|---|---|---|---|
| Acrosin | Pig | 40 - 283 | 1FIZ* | A | 16 - 238 | 2.90 | Yes |
| Acrosin | Sheep | 40 - 283 | 1FIW* | A | 16 - 238 | 2.10 | Yes |
| Azurocidin | Human | 27 - 239 | 1A7S* | A | 1 - 213 | 1.12 | No |
| Cathepsin G | Human | 21 - 238 | 1AU8 | A | 16 - 238 | 1.90 | Yes |
| Chymotrypsin-Like Elastase 1 | Pig | 27 - 259 | 1B0E* | A | 16 - 238 | 1.80 | Yes |
| Chymotrypsin-Like Elastase 2 | Pig | 29 - 262 | 1BRU | P | 16 - 238 | 2.30 | Yes |
| Complement Factor D | Human | 26 - 248 | 1BIO* | A | 16 - 238 | 1.50 | Yes |
| Complement Factor D | Mouse | 26 - 249 | 5FCR* | A | 16 - 238 | 1.25 | No |
| Complement Factor I | Human | 340 - 569 | 2XRC* | B | 322 - 551 | 2.69 | No |
| Complement Factor 1R | Human | 464 - 697 | 1GPZ* | A | 447 - 680 | 2.90 | No |
| Complement Factor 1S | Human | 438 - 675 | 1ELV | A | 423 - 660 | 1.70 | No |
| Chymase | Human | 22 - 240 | 1KLT* | A | 16 - 238 | 1.90 | Yes |
| Chymotrypsinogen B | Rat | 169 - 256 | 1KDQ* | B | 151 - 238 | 2.55 | No |
| Chymotrypsin-C | Bovine | 30 - 262 | 1PYT | D | 716 - 938 | 2.35 | No |
| Chymotrypsin-C | Human | 30 - 262 | 4H4F* | A | 16 - 238 | 1.90 | Yes |
| Duodenase-1 | Bovine | 20 - 238 | 1EUF* | A | 16 - 238 | 2.40 | No |
| Enteropeptidase | Bovine | 801 - 1030 | 1EKB | B | 16 - 238 | 2.30 | Yes |
| Coagulation Factor VII | Human | 213 - 447 | 1CVW* | H | 16 - 238 | 2.28 | Yes |
| Coagulation Factor IX | Human | 227 - 454 | 1RFN | A | 16 - 238 | 2.80 | Yes |
| Coagulation Factor IX | Pig | 183 - 409 | 1PFX | C | 16 - 237 | 3.00 | Yes |
| Coagulation Factor X | Bovine | 234 - 461 | 1KIG | H | 16 - 238 | 3.00 | Yes |
| Coagulation Factor X | Human | 235 - 462 | 1C5M* | D | 16 - 238 | 1.95 | No |
| Coagulation Factor XI | Human | 388 - 618 | 1XX9* | A | 16 - 238 | 2.20 | Yes |
| Coagulation Factor XII | Human | 373 - 609 | 6B74* | B | 16 - 238 | 2.32 | Yes |
| Coagulation Factor XIV | Human | 212 - 445 | 1AUT | C | 16 - 238 | 2.80 | Yes |
| Granzyme A | Human | 29 - 254 | 1OP8* | A | 16 - 238 | 2.50 | No |
| Granzyme B | Human | 21 - 240 | 1FQ3 | A | 16 - 238 | 3.10 | No |

| | | | | | | | |
|---|---|---|---|---|---|---|---|
| Granzyme B | Rat | 21 - 241 | 1FI8 | A | 16 - 238 | 2.20 | Yes |
| Granzyme C | Mouse | 21 - 241 | 3FZZ* | A | 21 - 241 | 2.50 | No |
| Granzyme H | Human | 21 - 239 | 3TJU* | A | 16 - 234 | 2.70 | Yes |
| Granzyme K | Human | 27 - 254 | 1MZA* | A | 16 - 238 | 2.23 | No |
| Granzyme M | Human | 26 - 249 | 2ZGC* | A | 1 - 224 | 1.96 | No |
| Haptoglobin | Human | 162 - 399 | 4WJG* | 2 | 158 - 395 | 3.10 | No |
| Haptoglobin | Pig | 103 - 340 | 4F4O* | C | 103 - 340 | 2.90 | No |
| Hepatocyte Growth Factor | Human | 495 - 716 | 1SHY* | A | 495 - 716 | 3.22 | No |
| Hepatocyte Growth Factor A | Human | 408 - 641 | 1YBW* | A | 408 - 641 | 2.70 | No |
| Hepatocyte Growth Factor L | Human | 484 - 704 | 2ASU* | B | 484 - 704 | 1.85 | No |
| Hepsin | Human | 163 - 400 | 1O5E* | H | 16 - 238 | 1.75 | Yes |
| Kallikrein-1 | Human | 25 - 254 | 1SPJ* | A | 16 - 238 | 1.70 | No |
| Kallikrein-2 | Horse | 25 - 253 | 1GVZ | A | 16 - 238 | 1.42 | No |
| Kallikrein-2 | Human | 25 - 253 | 4NFE* | A | 16 - 238 | 1.90 | Yes |
| Kallikrein-2 | Rat | 25 - 251 | 1TON | A | 16 - 238 | 1.80 | No |
| Kallikrein-3 | Human | 25 - 253 | 2ZCH* | P | 16 - 238 | 2.83 | No |
| Kallikrein-4 | Human | 31 - 247 | 2BDG* | A | 16 - 238 | 1.95 | Yes |
| Kallikrein-5 | Human | 67 - 285 | 2PSX* | A | 16 - 238 | 2.30 | Yes |
| Kallikrein-6 | Human | 22 - 237 | 1GVL | A | 16 - 238 | 1.80 | No |
| Kallikrein-7 | Human | 30 - 245 | 2QXG* | A | 16 - 238 | 2.60 | Yes |
| Kallikrein-8 | Human | 33 - 252 | 5MS3* | A | 16 - 238 | 2.30 | No |
| Kallikrein-8 | Mouse | 33 - 252 | 1NPM | A | 16 - 238 | 2.10 | No |
| Kallikrein-10 | Human | 50 - 269 | 5LPE* | B | 20 - 238 | 2.65 | No |
| Kallikrein-13 | Mouse | 25 - 253 | 1AO5* | A | 16 - 238 | 2.60 | No |
| Kallikrein-Related B1 (KLKB1) | Human | 391 - 621 | 2ANW* | A | 16 - 238 | 1.85 | Yes |

| | | | | | | | |
|---|---|---|---|---|---|---|---|
| Kallikrein-Related B1 (KLKB1) | Mouse | 391 - 621 | 5GVT* | A | 16 - 238 | 2.61 | No |
| Kallikrein-Related B3 (KLKB3) | Mouse | 25 - 253 | 1SGF* | G | 16 - 238 | 3.15 | No |
| Kallikrein-Related B4 (KLKB4) | Mouse | 29 - 248 | 1SGF* | A | 25 - 238 | 3.15 | No |
| Mast Cell Protease 2 | Rat | 21 - 239 | 3RP2* | A | 16 - 238 | 1.90 | No |
| MBL Serine Protease 1 | Human | 449 - 691 | 3GOV* | B | 449 - 691 | 2.55 | No |
| MBL Serine Protease 2 | Human | 445 - 679 | 1Q3X* | A | 445 - 679 | 2.23 | No |
| Membrane-Type Serine Protease 1 | Human | 615 - 849 | 1EAW | A | 16 - 238 | 2.93 | Yes |
| Myeloblastin | Human | 28 - 243 | 1FUJ* | A | 16 - 238 | 2.20 | No |
| Neutrophil Elastase | Human | 30 - 242 | 1B0F* | A | 16 - 238 | 3.00 | Yes |
| Proproteinase-E | Bovine | 12 - 246 | 1PYT | C | 416 - 638 | 2.35 | No |
| Prostatin | Human | 45 - 281 | 3DFJ* | A | 45 - 281 | 1.45 | No |
| Serine Protease 57 | Human | 34 - 258 | 4Q7X* | A | 16 - 238 | 2.55 | No |
| Thrombin B | Human | 364 - 430 | 2HNT | C | 16 - 72 | 2.50 | No |
| Thrombin B | Mouse | 361 - 610 | 2OCV* | B | 16 - 238 | 2.20 | No |
| Tissue Plasminogen | Human | 311 - 556 | 1A5H | A | 16 - 238 | 2.90 | Yes |
| Trypsin | Boar | 9-231 | 1AVW* | A | 9-231 | 1.75 | Yes |
| Trypsin | Pig | 134 - 224 | 1AKS* | B | 146 - 238 | 1.80 | No |
| Trypsin-1 | Bovine | 24 - 239 | 1AQ7* | A | 16 - 238 | 2.20 | Yes |
| Trypsin-2 | Rat | 24 - 239 | 1AMH* | A | 16 - 238 | 2.50 | No |
| Trypsin-3 | Human | 81 - 296 | 1H4W* | A | 16 - 238 | 1.70 | Yes |
| Trypsin B1 | Human | 31 - 267 | 1LTO | A | 16 - 238 | 2.20 | No |
| Trypsin B2 | Human | 31 - 267 | 1A0L | A | 16 - 238 | 3.00 | Yes |
| Urokinase | Human | 179 - 419 | 1C5W* | B | 16 - 238 | 1.94 | Yes |
| Urokinase | Mouse | 180 - 421 | 5LHN* | A | 16 - 238 | 2.55 | Yes |

**Supplementary Table 2. Structures used in the structure-based phylogenetics analysis**

** - denotes structural models re-refined in PDB-REDO.*

| Inhibitor(s) | Disease/Indication | Target(s) | Phase |
|---|---|---|---|
| Camostat mesylate | SARS-CoV-1<br>SARS-CoV-2 | TMPRSS2 | Preclinical |
| Nafamostat mesylate | MERS-CoV<br>SARS-CoV-2 | TMPRSS2 | Preclinical |
| Avoralstat | Angioedema | Plasma Kallikrein | Phase 3 |
| PCI-27483 | Pancreatic Cancer | Factor VII | Phase 2 |
| Antipain | Biochemical Assays | Trypsin/Papain | Experimental |
| Leupeptin | Biochemical Assays | Trypsin/Papain | Experimental |
| Hepsin Inhibitors (H1 – 12) | Prostate Cancer | Hepsin | Experimental |
| Trypsin Inhibitors (T1 – 6) | Biochemical Assays | Trypsin | Experimental |
| Factor VII Inhibitors (FVII1 – 19) | Coagulopathy | Factor VII | Experimental |
| Factor XI Inhibitors (FXI1 – 32) | Coagulopathy | Factor XI | Experimental |
| Acrosin Inhibitors (A1 – 16) | Contraception | Acrosin | Experimental |
| Plasma Kallikrein Inhibitors (K1 – 5) | Prostate Cancer | Plasma Kallikrein | Experimental |

**Supplementary Table 3. Inhibitors targeting TMPRSS2 and structurally similar proteases**

| Inhibitor | Structure | Docking Tool | S1-Peptidase Domain | | | |
|---|---|---|---|---|---|---|
| | | | TMPRSS2 | KLKB1 | Factor VII | Trypsin |
| SARS-CoV-2 Spike S2’ | $^{+}H_3N$-KPSKRSF-$COO^{-}$ | Glide | -9.67 | -10.68 | -8.56 | -10.02 |
| | | HADDOCK | -97.0 (2.7) | -113 (1.3) | -92.2 (2.2) | -79.3 (2.7) |
| Camostat | 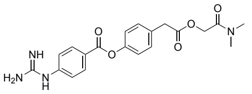 | Glide | -4.68 | 0.45 | -4.14 | N.D.* |
| | | HADDOCK | -51.6 (2.3) | -48.4 (2.0) | -42.4 (0.9) | -42.7 (0.2) |
| Avoralstat | 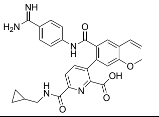 | Glide | -7.78 | -5.59 | -1.74 | -4.22 |
| | | HADDOCK | -70.6 (0.7) | -78.1 (2.6) | -56.0 (1.1) | -56.5 (0.5) |
| PCI-27483 | 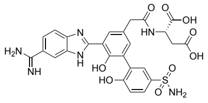 | Glide | -8.78 | -5.63 | -12.7 | -9.15 |
| | | HADDOCK | -106 (2.7) | -81.5 (1.2) | -84.7 (1.3) | -90.2 (5.2) |
| Antipain | 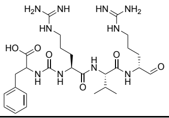 | Glide | -9.77 | -6.71 | -8.98 | -8.98 |
| | | HADDOCK | -75.1 (1.4) | -72.7 (4.4) | -61.2 (3.6) | -56.8 (2.2) |
| Leupeptin | 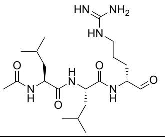 | Glide | -7.08 | -5.54 | -8.22 | -7.57 |
| | | HADDOCK | -49.6 (1.6) | -48.1 (2.6) | -35.3 (1.8) | -41.1 (2.6) |
| MDL-28170 | 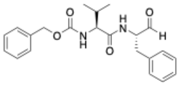 | Glide | -5.55 | -7.68 | -6.31 | -5.41 |
| | | HADDOCK | -39.2 (1.3) | -40.8 (3.2) | -29.0 (1.2) | -37.0 (0.7) |
| Ritonavir | 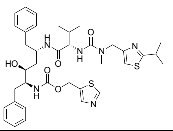 | Glide | -6.02 | -6.82 | -5.10 | -6.15 |
| | | HADDOCK | -53.5 (1.4) | -54.0 (1.6) | -44.4 (4.0) | -46.4 (2.6) |
| Lopinavir | 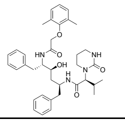 | Glide | -6.33 | -8.13 | -6.18 | -6.75 |
| | | HADDOCK | -45.4 (1.1) | -42.6 (3.1) | -43.5 (2.1) | -50.5 (1.3) |
| SBTI Peptide | $^{+}H_3N$-PWRIRF-$COO^{-}$ | Glide | -9.18 | -9.35 | -7.50 | -7.97 |
| | | HADDOCK | -77.3 (0.5) | -82.5 (1.3) | -58.2 (2.5) | -67.5 (2.5) |

**Supplementary Table 4. Docking scores of protease inhibitors and substrates to TMPRSS2-S1P and related peptidase domains.** * - denotes the compound did not result docking score. Numbers in parentheses indicates STDEV.

| S1-Peptidase Domain | TMPRSS2 | Trypsin | KLKB1 | Factor VII |
|---|---|---|---|---|
| **PDB ID** | Model | 1AVW | 6O1S | 1W7X |
| **HADDOCK Score (STDEV)** | -154.6 (7.5) | -171.7 (1.3) | -140.1 (6.1) | -87.7 (2.0) |
| **Z-Score** | -1.8 | -1.4 | -1.7 | -1.5 |

**Supplementary Table 5. Docking scores of SBTI to TMPRSS2-S1P and related S1-peptidase domains**

| Protease | Avoralstat | PCI-27483 | Antipain | SBTI | Camostat |
|---|---|---|---|---|---|
| **TMPRSS2** | 2.73 ± 0.19 nM | 1.41 ± 0.04 μM | 748 ± 63 nM | 121 ± 4 nM | 1.01 ± 0.10 nM |
| **KLKB1** | 127 ± 4 pM | N.I. | 718 ± 153 nM | 7.04 ± 0.51 nM | 446 ± 140 pM |
| **Trypsin** | 48.8 ± 0.3 nM | 2.90 ± 0.14 μM | 16.2 ± 5.1 nM | 204 ± 13 pM | 280 ± 84 pM |
| **Factor VIIa** | 122 ± 16 nM | 82.8 ± 4.9 nM | N.I. | N.I. | 4.76 ± 0.06 uM |
| **Factor Xa** | 196 ± 6 nM | 5.16 ± 0.06 μM | 2.21 ± 0.06 μM | 132 ± 10 nM | 8.87 ± 0.36 uM |
| **Papain** | N.I. | N.I. | 964 ± 185 pM | N.I. | N.I. |
| **KLK1** | 321 ± 23 nM | N.I. | N.I. | N.I. | 768 ± 208 nM |
| **KLK7** | N.I. | N.I. | N.I. | 61.5 ± 8.4 nM | N.I. |
| **Furin** | N.I. | N.I. | N.I. | N.I. | N.I. |
| **Mpro** | N.I. | N.I. | N.I. | N.I. | N.I. |
| **PLpro** | N.I. | N.I. | N.I. | N.I. | N.I. |

**Supplementary Table 6. $IC_{50}$ of the four 3DPhyloFold inhibitors and Camostat against ten proteases.** Six S1-peptidases identified in 3DPhyloFold (i.e., KLKB1, Trypsin, Factor VIIa, Factor Xa, KLK1, and KLK7), three other proteases involved in SARS-CoV-2 infection (i.e., Furin, Mpro, and PLpro), and a negative control Papain was tested. Data are represented as mean ± SEM (n = 3). N.I. indicates not inhibiting ($IC_{50}$ over determinable range).

| Protease | Avoralstat $IC_{50}$ (M) |
|---|---|
| **ACE1** | N.I. |
| **ACE2** | N.I. |
| **ADAM10** | 6.06E-05 |
| **BACE1** | N.I. |
| **Calpain 1** | N.I. |
| **Caspase 1** | 4.89E-06 |
| **Caspase 10** | N.I. |
| **Caspase 11** | 1.06E-06 |
| **Caspase 14** | 7.82E-06 |
| **Caspase 2** | 2.57E-06 |
| **Caspase 3** | 7.29E-06 |
| **Caspase 4** | N.I. |
| **Caspase 5** | N.I. |
| **Caspase 6** | 3.07E-07 |
| **Caspase 7** | 6.18E-06 |
| **Caspase 8** | 4.47E-07 |
| **Caspase 9** | N.I. |
| **Cathepsin B** | 5.34E-05 |
| **Cathepsin C** | 2.55E-05 |
| **Cathepsin D** | 1.87E-05 |

| Protease | Avoralstat $IC_{50}$ (M) |
|---|---|
| **Cathepsin E** | N.I. |
| **Cathepsin G** | 3.08E-06 |
| **Cathepsin H** | 2.23E-05 |
| **Cathepsin K** | 7.75E-06 |
| **Cathepsin L** | 4.87E-05 |
| **Cathepsin S** | 2.22E-05 |
| **Cathepsin V** | 3.50E-06 |
| **Chymase** | 2.98E-05 |
| **Chymotrypsin** | 1.02E-05 |
| **DPP IV** | 7.76E-05 |
| **DPP IX** | 2.03E-05 |
| **DPP VIII** | 2.51E-05 |
| **Elastase** | 4.00E-05 |
| **Factor XIa** | 1.91E-08 |
| **HIV-1** | N.I. |
| **HTRA1** | N.I. |
| **Kallikrein 12** | <3.81E-10 |
| **Kallikrein 13** | 3.33E-07 |
| **Kallikrein 14** | 5.10E-10 |
| **Kallikrein 5** | 1.44E-08 |

| Protease | Avoralstat $IC_{50}$ (M) |
|---|---|
| **Matriptase 2** | 2.73E-09 |
| **MMP 1** | N.I. |
| **MMP 10** | 5.68E-06 |
| **MMP 12** | 1.73E-05 |
| **MMP 13** | 1.33E-05 |
| **MMP 14** | 6.21E-05 |
| **MMP 2** | 1.72E-05 |
| **MMP 3** | 1.98E-05 |
| **MMP 7** | 2.28E-05 |
| **MMP 8** | 2.45E-05 |
| **MMP 9** | 1.70E-05 |
| **Neprilysin** | 3.28E-05 |
| **Plasmin** | 1.76E-09 |
| **Proteinase A** | N.I. |
| **Proteinase K** | N.I. |
| **TACE** | 2.11E-05 |
| **Thrombin a** | 2.85E-08 |
| **Tryptase b2** | 9.96E-10 |
| **Tryptase g1** | 9.29E-10 |
| **Urokinase** | 8.17E-08 |

**Supplementary Table 7. Specificity profile of Avoralstat against 60 proteases.** N.I. indicates not inhibiting ($IC_{50}$ over determinable range). <3.81E-10 indicates $IC_{50}$ below determinable range.
